# Multi-omics uncover interactions in the vaginal microbiome and a type II secretion/Tad pilus system in *Gardnerella* species

**DOI:** 10.64898/2026.05.09.724037

**Authors:** Fabricio Romero García, Vladyslav Dovhalyuk, Sen Lang Kuilboer, Karlijn Johanna van Dijk, Ye Liu, Clarissa Forsström, Hassan Gharibi, Atef Mahmoud Mannaa, Ákos Végvári, Anders Karlsson, Roger Karlsson, Per Hydbring, Luisa Warchavchik Hugerth, Cammie Lesser, Lars Engstrand, Amir Ata Saei, Daniel Globisch, Juan Du

## Abstract

**Background:** The vaginal microbiome is a key determinant of women’s health, with *Lactobacillus*-dominant communities protecting against bacterial vaginosis (*BV*) and related adverse outcomes. Dysbiosis involves a decline in lactobacilli and overgrowth of anaerobes, with *Gardnerella* species central to initiating polymicrobial aggregates. However, the genetic determinants of community assembly, biofilm formation, and interspecies interactions remain poorly understood. This study investigated the genomic and functional basis of these traits using an integrated multi-omics approach.

**Results:** Long-read whole-genome sequencing of clinical *Gardnerella* isolates yielded closed genomes and revealed a pangenomic split correlating with high-versus low-biofilm phenotypes, linked to clade-specific genomic islands and allelic variants. Functional gene modules within these regions distinguished *G. vaginalis* from *G. piotii*, contributing to nutrient acquisition, cell wall metabolism, and colonization. Integrated 16S rRNA gene sequencing, and proteomics of all pairwise dual-species co-cultures of five key vaginal bacteria showed that *G. vaginalis* and *L. iners* acted as dominant ecological drivers, outcompeting neighbors and triggering asymmetric proteomic remodeling. Differential expression of a gene within a putative operon encoding a conserved type II secretion/Tad pilus system, revealed by integrating the pangenome with the proteome, implicated this system in interspecies interactions. AI-based structural predictions of the complex suggested isoform-specific differences in protein folding between *G. vaginalis* and *G. piotii*. In parallel, metagenomic analysis of 93 women showed the operon was more abundant in *BV* samples than in healthy controls, linking it to dysbiosis. Integrated metabolomics further identified a methyl-β-carboline compound elevated in *Prevotella bivia*-containing co-cultures that selectively inhibited *G. vaginalis* and *P. bivia* while sparing *L. crispatus*. Several dipeptides influenced growth and biofilm formation of vaginal microbes, and methyl-β-carboline and sphingosine additionally inhibited multidrug-resistant pathogens, suggesting new strategies for managing polymicrobial infections.

**Conclusions:** This work links *Gardnerella* genomic features to virulence-associated phenotypes and demonstrates how interspecies interactions shape community dynamics and antimicrobial metabolite production. The findings support a model in which biofilm capacity is genetically encoded within an accessory genome, deployed through tailored proteomic responses, and modulated by an interaction-specific metabolic landscape, providing a foundation for identifying bioactive molecules and developing diagnostics and targeted therapeutics to restore and maintain vaginal health.

**Highlights:**

1. Long-read whole-genome sequencing of clinical *Gardnerella* isolates yielded closed ge-nomes and linked high-versus low-biofilm phenotypes to clade-specific genomic islands and allelic variants.
2. Integrated 16S rRNA gene sequencing, proteomics, and metabolomics of all pairwise du-al-species co-cultures showed that *Gardnerella vaginalis* and *Lactobacillus iners* outcompete neighboring species and trigger asymmetric proteomic remodeling, identifying them as dominant ecological drivers of dysbiosis.
3. AI-predicted structural variation in a conserved type II secretion/Tad pilus system, sup-ported by metagenomic cohort analysis of 93 women, distinguished healthy from dysbiotic states and highlights a candidate BV biomarker.
4. Integrated metabolomics identified a methyl-β-carboline elevated in *Prevotella bivia*-containing co-cultures that selectively inhibits *G. vaginalis* and *P. bivia* while sparing *L. crispatus*, offering a narrow-spectrum therapeutic lead.
5. Metabolites including several dipeptides modulated growth and biofilm of vaginal mi-crobes and multidrug-resistant pathogens, suggesting new strategies for managing polymicrobial infections

## Background

The human microbiome is a critical determinant of health and disease, with its composition and function influencing systemic physiology, immune responses, and pathogen resistance [1]. Within this ecosystem, the vaginal microbiome is particularly important for obstetric health [2]. A stable, low-diversity community dominated by *Lactobacillus* species confers protection through, for example, lactic acid production, maintenance of low pH, and competitive exclusion of pathogens [3]. Conversely, dysbiosis, often characterized by a decline in lactobacilli and an increase in diverse anaerobic bacteria, is linked to adverse outcomes including bacterial vaginosis (*BV*), increased risk of sexually transmitted infections, preterm birth, and other complications [4].

Among the key taxa associated with dysbiosis, *Gardnerella* species play a central role [5]. Although frequently detected in asymptomatic individuals [6], *Gardnerella vaginalis* (*Gv*) is recognized as the primary initiator of polymicrobial biofilms, with secondary contributors including *Prevotella bivia* (*Pb*) and *Fannyhessea vaginae* (*Fv*) [7],[8] [9]. However, the genomic and functional diversity within *Gardnerella* is not yet fully resolved [10], making it difficult to distinguish between commensal strains and variants driving biofilm formation [11],[12]. Recent revisions of the *Gardnerella* genus have led to the definition of additional genomic species, including *Gardnerella piotii* (*Gp*). These revisions continue to shape our understanding of the role of different *Gardnerella* species in disease etiology [13].

Furthermore, polymicrobial biofilms are clinically significant because they create a protective niche that enhances bacterial adherence, leading to chronic infections that pose risks to vaginal health [14]. Despite this understanding, the molecular cues mediating microbial-microbial interactions within the vaginal microbiome remain largely unknown [15]. These interspecies dynamics are driven by the exchange of secreted proteins and metabolites have yet to be characterized [16]. Additionally, bacterial secretion systems delivering potential virulence factors in the vaginal environment remain unexplored. Deciphering secretion and adhesion mechanisms is essential, as they likely serve as the functional link between *Gardnerella* genomic diversity and polymicrobial assembly [17],[18].

To address these knowledge gaps, we conducted an integrated multi-omics investigation of clinical vaginal isolates, focusing on *Gardnerella*. This included sequencing to create a high-quality genomic resource, phenotypic profiling of biofilm formation, and integrating dual-species co-culture models with downstream proteomics and metabolomics analyses. Our study revealed novel genetic elements and a conserved type II secretion/Tad pilus system (*T2SS/Tad*) in *Gardnerella*, together with pair-specific bioactive metabolites and proteins that correlate with biofilm phenotypes and may influence vaginal health (Fig. 1A).

**Figure 1.**
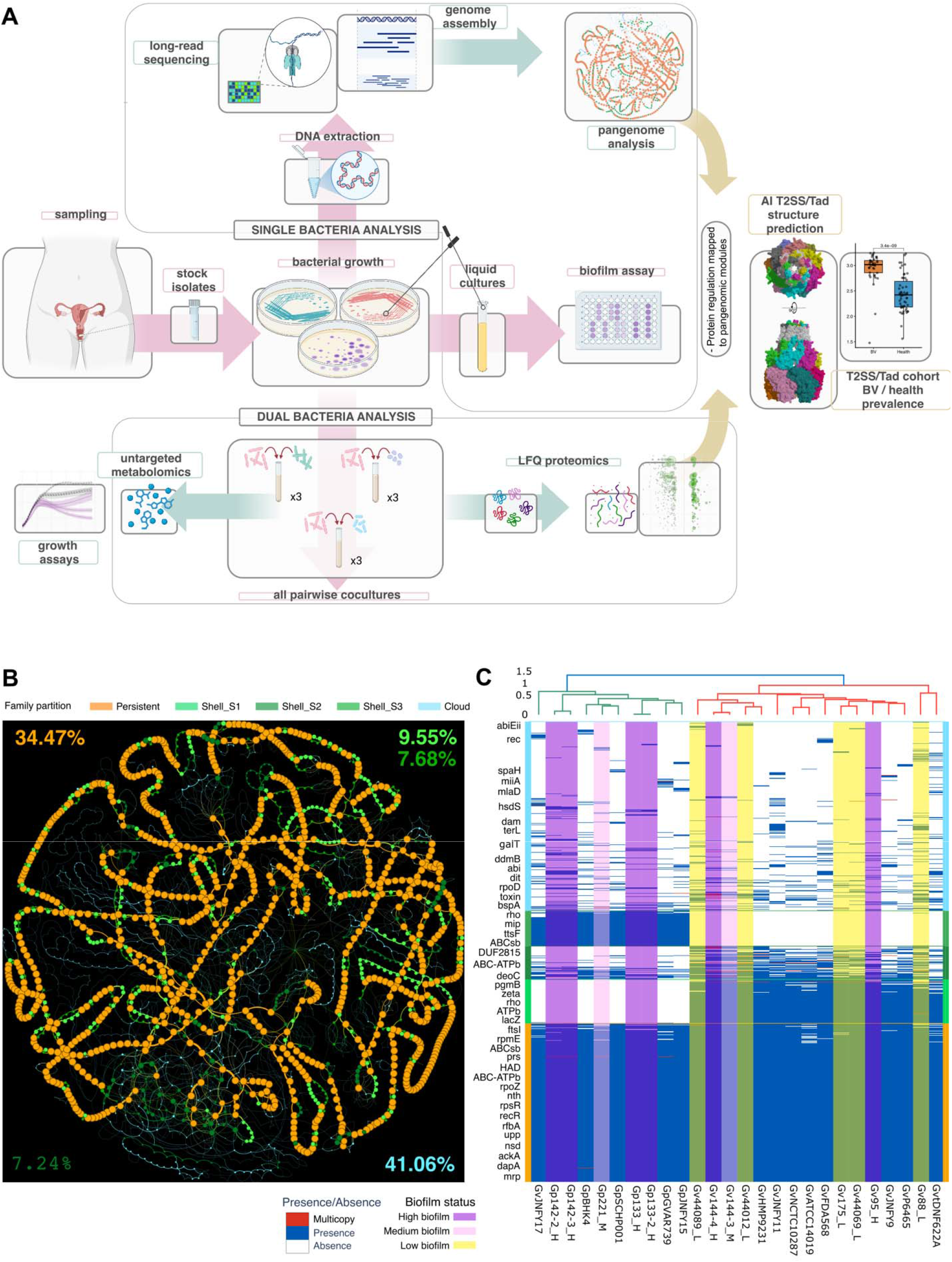
Accessory pangenome divergence links to biofilm capacity in *Gardnerella* species. A) Study workflow and experimental design. B) Syntenic network mapping gene families onto pangenome partitions in *Gardnerella*. Circles represent gene families, while edges represent genomic neighboring. Circle size represents the number of genomes in which the gene family is represented. Numbers in the corner represent the percentage of each partition in the pangenome, where: orange = persistent partition, light green = shell_S1 partition (*Gardnerella vaginalis*), dark green = shell_S2 partition (both or one *Gardnerella* species), medium green = shell_S3 partition (*Gardnerella piotii/pickettii*), light blue = cloud partition. C) Presence-absence matrix of gene families in *Gardnerella* genomes. Genomes are ordered by hierarchical clustering based on shared gene families using Jaccard distance. The gene families are ordered by partition and then by their number of presences (increasing order). Abbreviations BV = Bacterial vaginosis, *Gv* = *Gardnerella vaginalis*, *Gp* = *Gardnerella piotii/pickettii*. Full gene names for abbreviations used in the figure are provided in Supplementary Tables S3.

## Results

### Accessory pangenome divergence links to biofilm capacity in Gardnerella species

Vaginal biofilm formation supports *BV* persistence, with *Gardnerella* as a key colonizer [19]. Moreover, *Lactobacillus iners* (*Li*)*-*dominant communities are less stable and more prone to dysbiosis than those dominated by, for example, *L. crispatus* (*Lc*) [20]. Additionally, genomic determinants of biofilm formation in these species remain undefined, limiting understanding of disease mechanisms. To address these gaps, we undertook a comprehensive genomic and phenotypic analysis of *Gardnerella* and *Li*.

We first assessed biofilm formation across 15 *Gardnerella* isolates, comprising clinical and reference strains. Crystal violet staining revealed significant intra- and inter-species heterogeneity, with an overall mean OD_595_ of 0.61 ± 0.45. Based on this distribution, strains were categorized as high (mean OD_595_ [mOD_595_] > 0.76), medium (mOD_595_ 0.61 ± 0.15), or low (mOD_595_ < 0.46) biofilm formers. Biofilm biomass ranged from mOD_595_ 1.48 ± 0.20 for Gp133 to mOD_595_ 0.16 ± 0.07 for Gv88. Two isolates, Gp221 (mOD_595_ 0.52 ± 0.28) and Gv144-3 (mOD_595_ 0.53 ± 0.17), exhibited a medium biofilm-forming phenotype (Fig. S1a).

To investigate genomic architecture, we constructed a *Gardnerella* pangenome using 26 high-quality (> 99% completeness), closed circular genomes from our phenotyped isolates and public NCBI genomes (Tables S1a, b). The *Gardnerella* pangenome is partitioned into core (34.47%), accessory/shell (24.47%), and accessory/cloud (41.06%) gene families. Syntenic network analysis revealed that the core genomic backbone is frequently interrupted by accessory gene families, forming genomic islands or divergent gene families distributed across shell partitions or hypervariable loci (Fig. 1B). U-shaped and rarefaction plots, together with pangenome parameters support this conclusion and indicate an open accessory/cloud pangenome (Fig. S2b, c, and Table S2a). Hierarchical cluster analysis based on gene family presence/absence revealed a genomic split within the analyzed *Gardnerella* genome collection, separating *Gv* from *Gp*. This divergence correlated with a higher proportion of high-biofilm strains in *Gp* than in *Gv*, where each clade was defined by distinct genomic regions of conserved presence or absence (Fig. 1C). Together, these findings establish a pangenomic framework that links *Gardnerella* genomic diversity to *in vitro* biofilm-forming potential, revealing lineage-specific features that may underlie ecological insights into potential gene functions.

A similar analysis was performed on 11 *Li* strains, revealing that unlike *Gardnerella*, the *Li* genomes clustered according to a distinct pattern that did not show a comparable association with biofilm phenotype (Supplementary Information S2).

### Functional gene modules linked to clade-specific biofilm capacity in Gardnerella species

Analysis of genomic islands and divergent regions in *Gardnerella* revealed that they are primarily composed of modules of conserved gene families inserted at conserved *loci* across the analyzed genomes. These modules exhibited structural heterogeneity: some maintained synteny, whereas others were interspersed with core gene families; their size varied from 3 to 34 families. Crucially, many *loci* harbored distinct *alleles*, one enriched in *Gp* and another in *Gv*, differing in sequence identity and, less frequently, in gene family composition (Fig. S2a; Tables S4a, b).

Genes within modules of conserved gene families exhibited significantly higher sequence divergence between *Gp* and *Gv* than within either species, as determined by PPanGGOLiN. Several genomic islands were found exclusively in one species. Islands unique to *Gv* included a module for the galactose/Leloir pathway and peptidoglycan synthesis (Fig. S2b). Islands unique to *Gp* included a module for carbohydrate and micronutrient transport (Fig. 2A) and a toxin-antitoxin system (Fig. S2c). Notably, the Type I-E CRISPR-Cas system was present only in a subset of low-biofilm *Gv* strains. Interestingly, CRISPR-array sequences varied in length between strains (Fig. S2d). Other conserved modules, present in both species, diverged genetically: a cell wall metabolism module, with sequence identities ranging from 64.9 to 76.0%, was structurally conserved as a discrete island in *Gp* but fragmented in *Gv* by the insertion of three cell wall-anchoring and pilus-related proteins (Fig. 2B). Hypervariable gene hotspots included regions enriched in repeated motifs, including SpaH/EbpB family LPXTG-anchored major pilin, hypothetical proteins with heptad-repeat coiled-coil domains, peptidases, and FIVAR domain-containing proteins (Fig. 2C, D and S2e).

**Figure 2.**
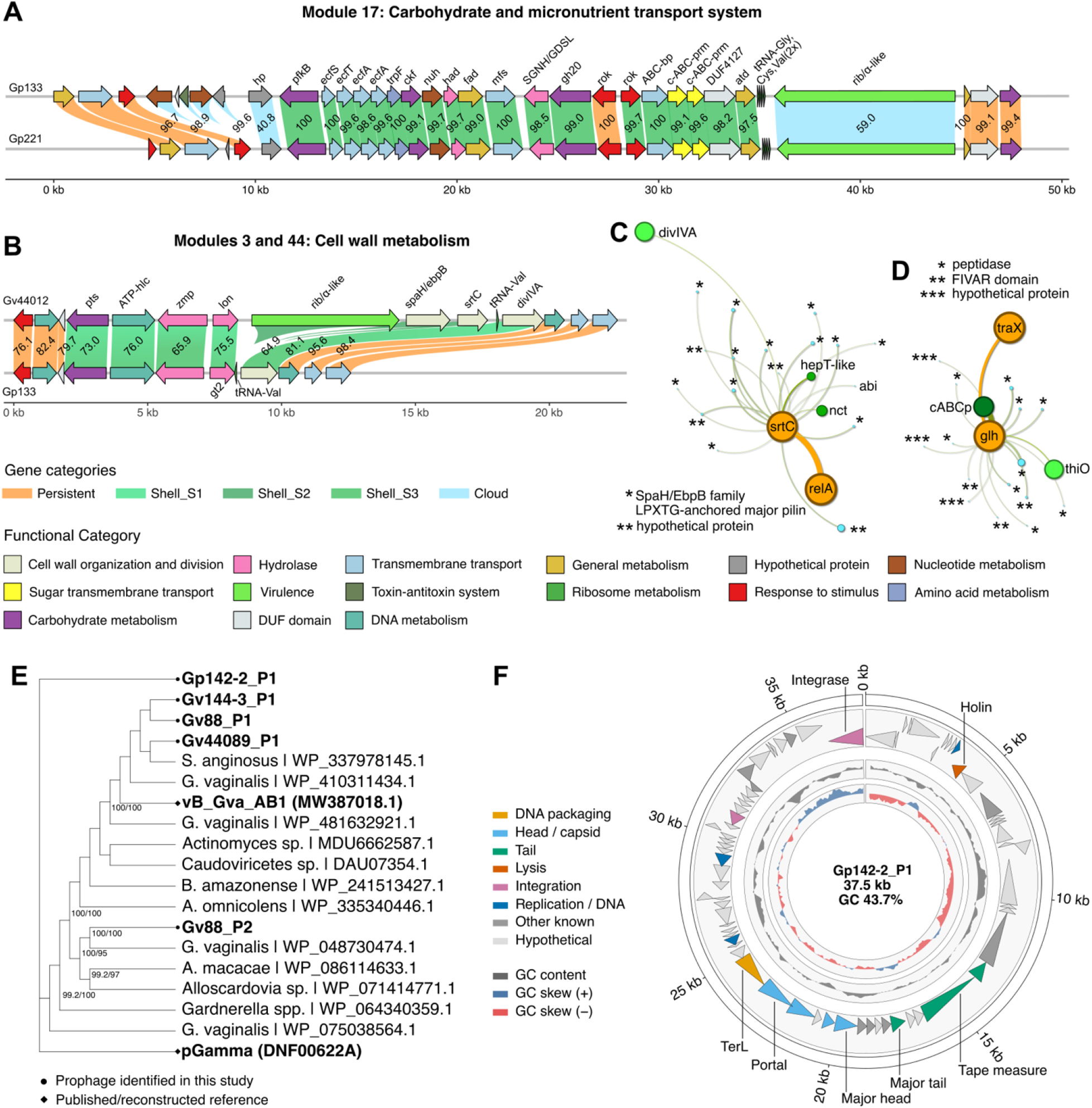
Functional gene modules linked to specific clades in *Gardnerella* species. Gene maps of relevant genomic modules. A) Module 17: Carbohydrate and micronutrient transport system. B) Modules 3 and 44: Cell wall metabolism. For additional genomic context, each module is depicted with three non-module gene families flanking its upstream and downstream regions. The vertical linking regions between synteny maps are coloured by pangenomic partition. Percentage amino acid identity between homologs is shown above each linking area. Network diagrams depict hypervariable gene families converging on core gene families, with the latter serving as gravitational centres. Circles represent gene families, while edges denote genomic neighbourhood relationships. Size of the circles represent the number of genomes the gene family is represented in; C) SpaH/EbpB family LPXTG-anchored major pilin (*) and hypothetical protein (**) converge on class C sortase (srtC), D) peptidase (*), FIVAR domain-containing protein (**), and hypothetical protein (***) converge on a glycoside hydrolase family protei (glh). E) TerL phylogeny of candidate prophages. Maximum-likelihood cladogram of 19 TerL amino-acid sequences comprising the five candidate prophages, the vB_Gva_AB1 and pGamma references, and 12 public homologues. Branch support assessed by 1,000 ultrafast bootstrap and 1,000 SH-aLRT replicates; node support is shown only where both values were ≥80%. Branch lengths are not proportional to evolutionary distance. F) Gv142-2_P1 prophage genome organization. Circular map of the 37.5-kb Gv142-2_P1 prophage (GC 43.7%), with CDSs coloured by functional category and oriented by transcription direction; key phage genes are labelled. Inner tracks show GC content and GC skew (500-bp window, 100-bp step). The circular layout is for visualization only and does not indicate circular topology. Abbreviations: *Gv* = *Gardnerella vaginalis*, *Gp* = *Gardnerella piotii/pickettii*. Full gene names for abbreviations used in the figure are provided in Supplementary Tables S3.

Moreover, an independent assessment of prophage sequences revealed that two strains, Gv88 and Gp142-2, each harbor a complete prophage. Notably, phylogenetic analysis based on the *terL* gene placed these prophages far apart (Fig. 2E). Beyond the essential phage machinery, which we assessed in the *Gp* prophage, the remaining proteins have yet to be characterized (Fig. 2F). Collectively, these findings show that species-specific genomic islands and the divergence of shared modules shape the genetic architecture underlying *in vitro* biofilm phenotypes in *Gardnerella*. In addition, phage elements, including complete prophages, are common in phylogenetically diverse *Gardnerella* strains.

### Species-specific proteomic interactions shape the dynamics of vaginal microbial co-cultures

Having characterized monospecies biofilm formation, we next modeled the vaginal microenvironment by examining interactions between common bacterial neighbors [21],[22]. We measured relative abundance changes and quantified bacterial protein expression in 24-hour all-pairwise co-cultures of five key vaginal bacteria. We defined the proteomic shift index (*PSI*) as the product of the proportion of significantly differentially abundant proteins (up- or downregulated; Welch’s t-test, *p* < 0.05) and their median fold change. *PSI* profiling revealed distinct, species-specific response patterns largely independent of relative abundance. *Gv* exhibited a moderate proteomic up-shift (mean up index [*MUI*] 22) and a pronounced down-shift (mean down index [*MDI*] -82), while outcompeting all co-culture partners (relative abundance [*RA*] 61.6-73.9%). *Li* displayed context-dependent remodeling, with a proteomic down-shift from modest with *Lc* (-27) to severe with *Gv* (-89), and its strongest up-shift with *Fv* (42), while representing the dominant member in most pairings (*RA* 38.4-75.0%). *Lc* exhibited a narrow proteomic shift across all partnerships (*MUI* 21, *MDI* -48), irrespective of dominance (*RA* 21.6-56.3%), suggesting comparatively moderate regulation. *Fv* displayed the most variable proteomic shifts, from an asymmetric response with *Lc* (up 7, down -73) to a nearly balanced bidirectional response with *Prevotella bivia* (up 65, down -63). *Pb*, by contrast, exhibited the second most variable proteomic response (*MUI* 22), with a downshift from -49 with *Fv* to -126 with *Li*, while being invariably outcompeted (*RA* 20.6-43.7%), indicating partner-driven proteomic remodeling decoupled from abundance (Fig. 3A). These data suggest partner-specific reprogramming, with *Gv* showing bidirectional remodeling and other species exhibiting changes weakly coupled to abundance.

**Figure 3.**
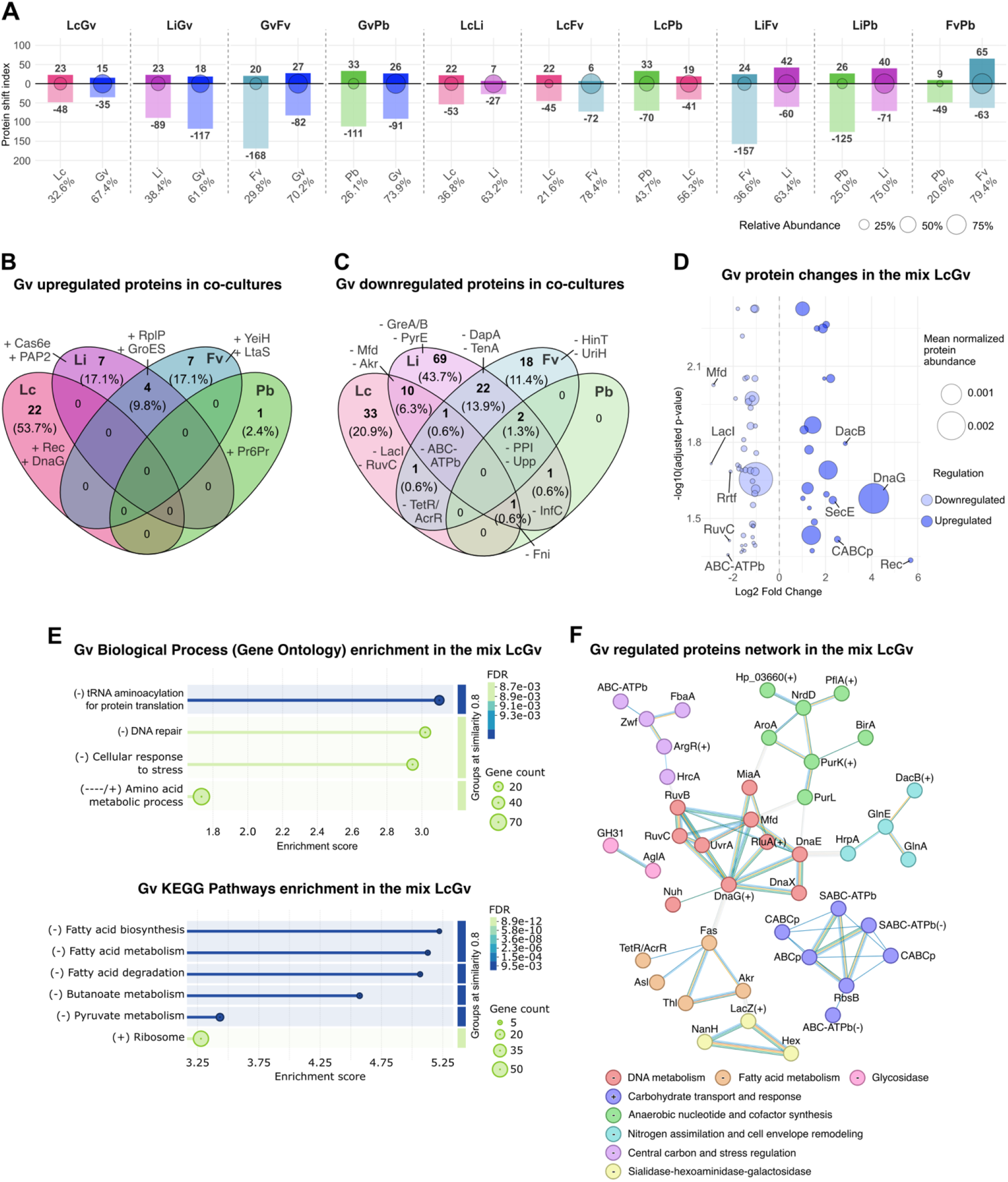
Species-specific proteomic interactions shape the dynamics of vaginal microbial co-cultures. A) Protein regulation and bacterial relative abundance in a co-culture system of all pairwise combinations of five different commensal vaginal bacteria. Protein shift index (PSI) per strain/condition is shown in the bar plots, and circle siz indicates 16S rRNA bacterial relative abundance. All conditions n=3, except for GvPb and FvPb, which are n=2. B) Venn diagram showing the number and relative percentage (based on the total number of regulated proteins) of upregulated proteins in *Gardnerella vaginalis (Gv)* when co-cultured with *Lactobacillus crispatus (Lc)*, *Lactobacillus iners (Li), Fannyhessea vaginae (Fv), and Prevotella bivia (Pb)*. The top two proteins are labeled per condition/overlap, when available. C) Venn diagram showing the number and relative percentage of downregulated proteins in Gv when co-cultured with *Lc, Li, Fv* and *Pb*. D) Bubble plot showing *Gv* protein changes in the co-culture *LcGv*. Downregulated proteins are on the left side in light colours and upregulated proteins on the right side. Circle size indicates the mean normalized protein abundance. The top 10 most fold-changed proteins are labelled, (FC > 1) adjusted *p-value* (< 0.05) (Welch’s t-test followed by a Benjamin-Hochberg multiple test correction). Strain abbreviations: *Lc = Lactobacillus crispatus, Li = Lactobacillus iners, Gv = Gardnerella vaginalis, Fv* = *Fannyhessea vaginae and Pb = Prevotella bivia.* E) Biological process (Gene Ontology) and KEGG pathway enrichment of Gv proteins during coculture with Lc. The x-axis shows the enrichment score; dot size represents the number of proteins annotated to each term; dot color indicates the false discovery rate (FDR), from dark blue (high significance) to pale green (lower significance). Only pathways passing the 1% FDR threshold are displayed. The sign before each term indicates: (-) = downregulation and (+) upregulation. F) Functional and physical protein association network of significantly changed Gv proteins during coculture with Lc. Nodes represent individual proteins; edges represent known or predicted functional associations, with line colour indicating the type of evidence (e.g., experimental, database, co-expression). The sign on top each group annotation node indicates: (-) = downregulation and (+) upregulation, while signs in individual proteins indicate exceptions in the group. Full protein names for abbreviations used in the figure are provided in Supplementary Tables S3.

To identify shared and condition-specific protein expression changes (|FC| > 1.0, adjusted *p* < 0.05), we compared the expression of 4,650 proteins identified across all co-culture pairs. *Gv* proteins regulated in response to other bacteria included the ribosomal assembly protein RplP, which was consistently upregulated with *Li* (log₂FC = 2.38) and *Fv* (2.51). Specific upregulation of Rec (DNA recombination, 5.69), Cas6e (adaptive immunity, 2.45), YeiH (sulfite sensitivity, 2.65), and Pr6Pr (membrane associated, 4.27) occurred with *Lc, Li, Fv,* and *Pb,* respectively (Fig. 3B, D and Fig. S3d-f). *Gv* proteins commonly downregulated included a PPIase (protein folding) with *Li* (-3.69)*, Fv* (-2.99), and *Pb* (-3.28); Mfd (transcription repair) with *Lc* (-2.85) and *Li* (-2.52); DapA (lysine biosynthesis) with *Li* (-5.43) and *Fv* (-4.09); and InfC (translation initiation) with *Li* (-1.34) and *Pb* (-1.00; Fig. 3C, D, and Fig. S3d-f). These data suggest that *Gv* intensifies defensive responses while downregulating transcriptional and protein-synthesis functions.

Assessing the *Gv* protein response to *Lc* is particularly relevant given the opposing roles these species play in dysbiosis and eubiosis, and their reciprocal interaction. Protein enrichment analysis of this *Gv* interaction revealed downregulation of tRNA aminoacylation for protein translation, DNA repair, cellular stress response, and amino acid metabolism (GO Biological Process), while KEGG analysis showed downregulated fatty acid, butanoate, and pyruvate metabolism and upregulated ribosomal pathways (Fig.3E). Network analysis revealed eight clusters of functionally related regulated proteins, the largest of which were involved in DNA metabolism, carbohydrate transport and response, and anaerobic nucleotide and cofactor synthesis (Fig.3F). Collectively, these findings reveal that *Gv* reprograms its translational and metabolic landscape in response to *Lc*, mounting a coordinated adaptive strategy that reshapes its physiology.

When co-cultured with *Gv*, other vaginal bacteria showed minimal common upregulation, limited to ATP-bc (membrane transport) in *Lc* (1.67) and *Li* (1.79). Species-specific responses were: *Lc* upregulated PriA (DNA replication, 5.35); *Li* increased PfkB (fructose metabolism, 2.76); *Fv* activated GlyT (glycoconjugate biosynthesis, 5.91); and *Pb* induced FtsE (cell division, 1.31) (Fig. S3a, c-f). Common downregulation targeted Prs (nucleotide production) in *Li* (-1.34) and *Fv* (-3.47), Dcd (pyrimidine metabolism) in *Lc* (-2.75) and *Li* (-2.41), and C1-Pep (peptide metabolism) in *Lc* (-1.59) and *Fv* (-2.58); Fig. S3b-f). Together, these analyses delineate a response to *Gv* characterized by conserved membrane transport upregulation and downregulation of nucleotide and protein metabolism, revealing partner-specific perturbations that rewire central metabolism, cellular homeostasis, and niche-specific functions in each co-cultured species. Additional proteomic analyses of *Lc* and *Li* co-cultures are presented in Supplementary Information S3.

### A genomic basis for co-culture-associated proteome shifts in G. vaginalis

To integrate our pangenome and proteome findings, we compared conserved gene modules linked to species divergence against protein expression changes mapped to these genomic regions. This analysis revealed 48 gene products, identified across all *Gv* co-culture mixes, that were linked exclusively to conserved modules in the *Gardnerella* pangenome (Tables S5a-e). These modules encompassed a broad spectrum of functions, including sugar uptake and utilization, cell wall dynamics, adherence, adaptive immunity, isoprenoid biosynthesis, and transcriptional regulation (Fig. 4A, B). Modules partitioned into two opposing groups: a smaller upregulated set, dominated by *Lc*-associated changes and enriched in membrane transport, carbohydrate utilization, and cell division functions, and a larger downregulated set, predominantly *Li*-associated and encompassing diverse metabolic and regulatory functions. Several modules showed opposing directionality across conditions, indicating that regulation is condition-dependent; for example, beta-galactosidase (LacZ) within module 54 (chromosome partitioning and fucose/galactose utilization) were upregulated with *Lc* but downregulated with *Fv* and *Li*. Notably, in response to *Li*, type I-E CRISPR-Cas proteins were regulated in opposite directions, with Cas6e (module 91) upregulated, while CasA (module 90) was downregulated (Fig. S4a). Collectively, the module-level organization reveals a coordinated and largely opposing rewiring of transport, carbohydrate metabolism, and transcriptional regulation between conditions.

**Figure 4.**
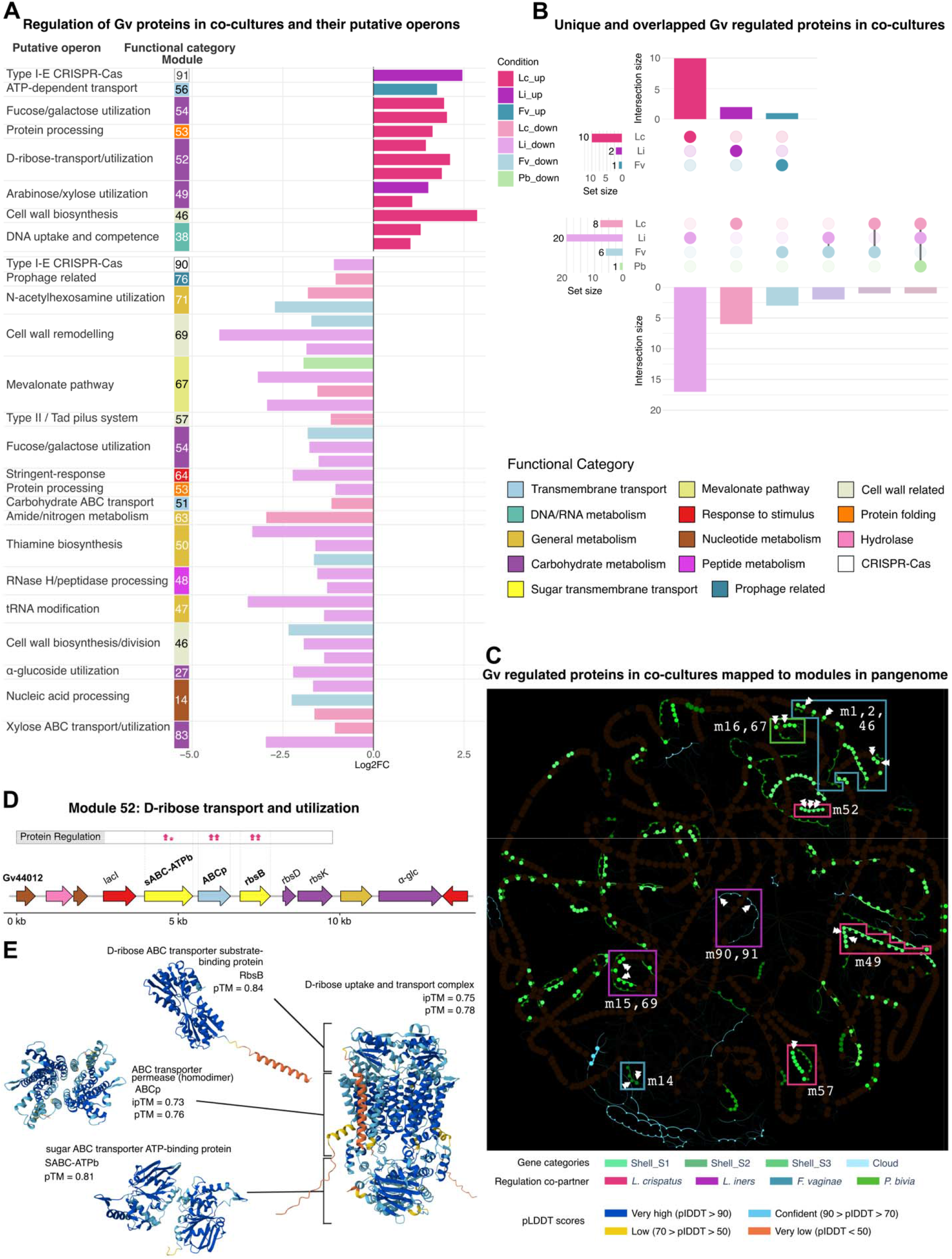
A genomic basis for co-culture-associated proteome shifts in *G. vaginalis*. Module-level regulation of Gv proteins across conditions. A) Divergent horizontal barplot showing log2 fold changes (adj. *p* < 0.05) of regulated proteins, grouped by module (putative operon) and coloured by condition. Modules are ordered by directionality, with a upregulated set (top) and a downregulated set (bottom). Putative operons are coloured by functional category. B) UpSet plots of down- and upregulated Gv proteins across conditions. UpSet plots showing intersections of regulated proteins between conditions for the upregulated (top) and downregulated (bottom) protein sets. Bars indicate intersection size, and the set-size bar plot (left) shows the total number of regulated proteins per condition (Lc, Li, Fv, Pb). Filled coloured dots denote the conditions participating in each intersection. Bars and dots are coloured by condition as in a. C) Syntenic network of conserved modules present in the accessory genome. Highlighted in colours according pangenome partition. Enclosed are genomic modules shown in a and in gene maps, where m## stands for the module number. White arrows indicate the specific gene families regulated in each module. Envelope colour indicates the main or some of the bacteria driving the regulation. Gene map; D) Module 52: Transcription-regulated D-ribose transport and metabolism. E) AI predicted protein structures of Gv’s D-ribose ABC transporter substrate-binding protein, pTM = 0.84; ABC transporter permease (homodimer), ipTM = 0.73, pTM = 0.76; sugar ABC transporter ATP-binding protein, pTM = 0.81; D-ribose uptake and transport complex, ipTM = 0.75, pTM = 0.78. For additional genomic context, each module is depicted with three non-module gene families flanking its upstream and downstream regions. Protein regulation: Regulated gene names are presented in bold with arrows indicating direction (one arrow = 1.0-1.4 |FC|; one and a half arrows = 1.45-1.85 |FC|, etc). Arrow colors indicate the bacterial species triggering the regulation, with same colour coding as C. Abbreviations: *Lc = Lactobacillus crispatus, Gv = Gardnerella vaginalis, Gp = Gardnerella piotii/pickettii, Li = Lactobacillus iners, Fv = Fannyhessea vaginae and Pb = Prevotella bivia*. Full gene/protein names for abbreviations used in the figure are provided in Supplementary Tables S3.

Closer inspection of these modules revealed a coherent set of genes. Overexpression of a sugar ABC transporter, an ABC permease, and RbsB, triggered by *Lc,* mapped to a D-ribose transport and utilization module (Fig. 4C, D). AI-predicted multimer models using the three upregulated protein sequences suggested a D-ribose uptake complex [23],[24] (Fig. 4E). Reduced expression of RecG triggered by *Li* and of Fni by *Lc*, *Li* and *Pb*, mapped to a module containing genes of a partial mevalonate pathway for isoprenoid biosynthesis. Module variation between species included species-specific distinctions in AI-predicted protein structures (Fig. 4C, and S4b). Upregulation of DacB induced by *Lc*, alongside downregulation of TilS by *Fv* and of MraZ and Mur by *Li*, mapped to a carbohydrate-responsive cell wall biosynthesis and division module. AI-based structure prediction of the transcriptional repressor MraZ revealed a canonical homo-octameric ring [25] (Fig. 4C, and S4c). Downregulation of HAD-IIB triggered by *Li* and of Lcp by *Li* and *Fv,* mapped to a module involved in DNA uptake and competence-associated cell wall remodeling. In *Gp*, the module lacks a metal ABC transporter, which may suggest species-specific differences in metal homeostasis (Fig. 4C, and S4d). Furthermore, TetR/AcrR showed reduced expression triggered by *Lc* and *Fv,* and Exo by *Li*; both mapped to a narrow module flanked by variable gene regions and non-coding regions present in some members of both *Gardnerella* species (Fig. 4C, and S4e). Together, these data reveal that co-culture partners influence the expression of *Gv* proteins encoded within genomic modules governing carbohydrate utilization, cell wall biosynthesis, competence, and adaptive immunity. This context-dependent proteomic remodeling, enabled by modular genome architecture, allows *Gv* to employ distinct functional programs tailored to each interacting species.

### AI-predicted structural divergence in a conserved type II secretion/Tad pilus complex underlies species variation in Gardnerella

We next focused on a putative operon containing genes for Tad pilus biosynthesis coupled to a type II secretion system (*T2SS/Tad*), consistently found in all *Gardnerella*, with differential expression of a neighboring DnaX implicating this system (Fig. S5a). Notably, the divergent region of the operon showed higher conservation among *Gv* strains (94.9 - 97.7% identity) than between *Gv* and *Gp* (57.7 - 73.7%), with the exception of the ATPase monomer TadA, which showed high conservation between species (82.7%) and was categorized as part of the core genome (Fig. 5A). To test the hypothesis that sequence divergence between species underlies variation in protein structure, we employed AI-based prediction. Analysis of functional domains in TadA revealed a nearly identical ATP-binding site with a conserved six-amino-acid segment (Ser-Gly-Lys-Thr-Thr-Leu). The TadA hexamer interface residues were also conserved, differing by a single aromatic residue: His to Tyr. Given the distinct chemical properties of these residues, this substitution likely alters inter-subunit contacts, suggesting differences in quaternary assembly or conformational changes [26] (Fig. 5B).

**Figure 5.**
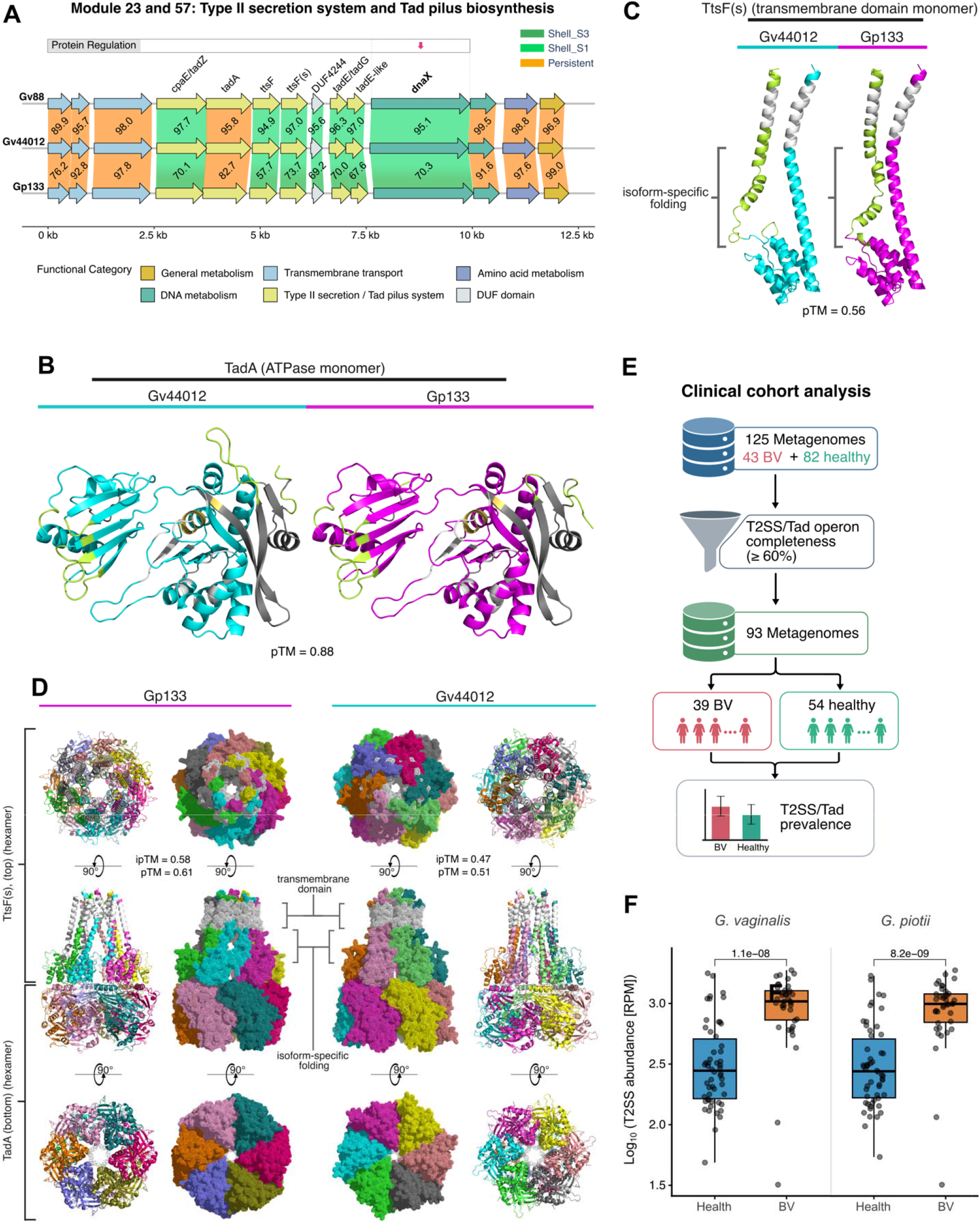
AI-predicted structural divergence in a conserved type II secretion/Tad pilus complex underlies species variation in *Gardnerella*. A) Gene map of type II secretion system/Tad pilus biosynthesis module. For additional genomic context, the module is depicted with three non-module gene families flanking its upstream an downstream region. Protein regulation: Regulated gene name is presented in bold with an arrow indicating direction (one arrow=1.0-1.4 |FC|). Arrow colour indicates the bacterial species triggering the regulation, where: dark-pink=*Lactobacillus crispatus*. The vertical linking regions between synteny maps are coloured by pangenomic partition. Percentage amino acid identity between homologs is shown on the linking area. Full gene/protein names for abbreviations used in the figure are provided in Extended Data Table S3. B) AI-predicted protein structure of TadA monomers. Cyan shows the main P-loop NTPase domain structure of Gv44012 (low biofilm strain), while magenta shows the structure of Gp133 (high biofilm strain). Yellow shows the ATP-binding domains; white marks regions of the hexamer interface; green highlights regions whose quaternary assembly is predicted to differ between species; and grey indicates the area outside the main ATPase structure. *pTM*=0.88, both structures. C) AI-predicted structure of TtsF(s) monomers. Colour coded as b. *pTM*=0.56 for both structures. D) AI-predicted architecture of TadA and TtsF(s) homohexamer complex. Cartoon and dotted surface representations show different details of the same angles. Subunits are coloured differentially, while the predicted transmembrane regions are coloured in white. Gp133 *ipTM*=0.58, *pTM*=0.61; Gv44012 *ipTM*=0.47; *pTM*=0.51. E) Workflow of T2SS/Tad clinical cohort analysis. F) Log_10_ completeness-adjusted abundance in reads per million (RPM) of the *T2SS/Tad* putative operon in *BV* and healthy metagenomic samples in both *Gardnerella* species. Differences between *BV* and healthy groups were assessed using the Wilcoxon rank-sum test. A *p-value* < 0.05 was considered statistically significant. Abbreviations: *BV = bacterial vaginosis, Gv = Gardnerella vaginalis, Gp = Gardnerella piotii, ipTM = interface predicted template modelling score, pTM = predicted template modelling score*.

We next assessed the structure of the type II secretion system F family protein (s) (*TtsF[s]*) monomer. Three regions, including a six-amino-acid segment at the *N*-terminus, a 36-amino-acid region adjacent to and partially overlapping the transmembrane domain, and a five-amino-acid segment at positions 120-124, may exhibit intrinsic flexibility or isoform-specific folding (Fig. 5C). To elucidate the molecular architecture of the putative secretion complex, we predicted the structure using the two described subunits, whose domains suggested a functional unit. The TadA complex formed a central homohexameric ring, an architecture canonical for secretion system ATPases, and potentially providing the energy for pilus polymerization [27],[28],[26]. Sharing interfaces with TadA, the TtsF(s) homohexamer containing predicted transmembrane alpha-helices may act as a structural coupler between the cytoplasmic ATPase and membrane-embedded components. Notably, prediction confidence was moderate for Gp133 (*ipTM* = 0.58, *pTM* = 0.61) and lower for Gv44012 (*ipTM* = 0.47, *pTM* = 0.51), warranting cautious interpretation. Despite these confidence limits, structural differences correlating with non-conservative amino acid substitutions suggest differences in complex stability, ATPase activity, or interactions with downstream Tad pilus subunits, which may contribute to observed phenotypic divergence (Fig. 5D).

We further explored the *T2SS/Tad* putative operon in clinical cohort data. Using public datasets, we evaluated *T2SS/Tad* completeness and abundance across 93 metagenomes (39 *BV*, 54 healthy controls; Fig. 5E) and found high prevalence in both *Gp* and *Gv* across all individuals, with higher abundance in *BV* samples (3.01 log_10_ mRPM [median reads per million]) than in healthy controls (2.44 log_10_ mRPM) (Wilcoxon rank-sum test, *p* < 0.001; Fig. 5F and S5b, c). These results indicate that genes within the putative operon are strongly associated with *BV*, likely due to the high prevalence of *Gardnerella* species. This suggests a disease-associated role for *Gardnerella*’s *T2SS/Tad* and raises the possibility of distinguishing *BV* from healthy conditions using a small protein panel.

### Untargeted metabolomics reveal dipeptides and other metabolites changed in co-cultures

Beyond protein-mediated interactions, microbial metabolites are key mediators of interbacterial competition and defense. To develop a more comprehensive model of the vaginal microenvironment, we identified shifts in relative metabolite abundance in supernatants from the same co-culture system (Fig. 6A) using a recently established workflow [29]. Species composition explained the largest share of metabolite variance between samples (46%): samples containing *Lactobacillus* clustered separately from those containing *Gv*, whereas samples containing only *Pb* or *Fv* were more dispersed (Fig. 6B). We then analyzed *Gv*-containing co-cultures for metabolites consistently altered in the opposite direction relative to monoculture controls. Significantly higher amounts of several dipeptides, including alanylproline (*p.adj* = 0.038-2×10^-4^, *t* = -2.766 to -14.17), lysylproline, phenylalanylproline, tyrosylproline, valylproline, glycylproline, and histidylproline, were detected in co-cultures containing *Gv* than in corresponding monocultures (Fig. 6A, C, and S6a). Notably, the natural abundance of these dipeptides, along with glycylmethionine, in *Lc* monoculture is linked to protein regulation in *LcGv* and *LcPb* co-cultures, where the dipeptidases PepDA (*p.adj* = 0.046, log_2_FC = -1.914) with *Gv* and PepDA (*p.adj* = 0.046, log_2_FC = 1.128) and PepX (*p.adj* = 0.049, log_2_FC = 3.561) with *Pb*, were significantly changed (Fig. 6D). Furthermore, the tripeptide valyl-prolyl-proline, which has a similar proline-based composition, was elevated in the *GvFv* co-culture compared to both monocultures (Fv, *p.adj* = 0.008, *t* = -3.748; Gv, *p.adj* = 2×10^-5^, *t* = - 17.32) (Fig. 6A, E). Using the same approach, we next searched for metabolites in *Pb* containing co-cultures. Three metabolites, tetrahydro-*β*-carboline-carboxylic acid, aspartylphenylalanine, and methyl-*β*-carboline (*p* = 0.001-5×10^-6^, *t* = -4.425 to 9.149), were detected at significantly higher levels in all *Pb*-containing co-cultures than in *Pb* monoculture, while sphingosine (*p.adj* = 0.011-7×10^-4^, *t* = 3.461 to -8.684) and urocanic acid were significantly elevated in all mono- and co-cultures containing *Pb* compared to co-partners monocultures (Fig. 6A, F, and S6b).

**Figure 6.**
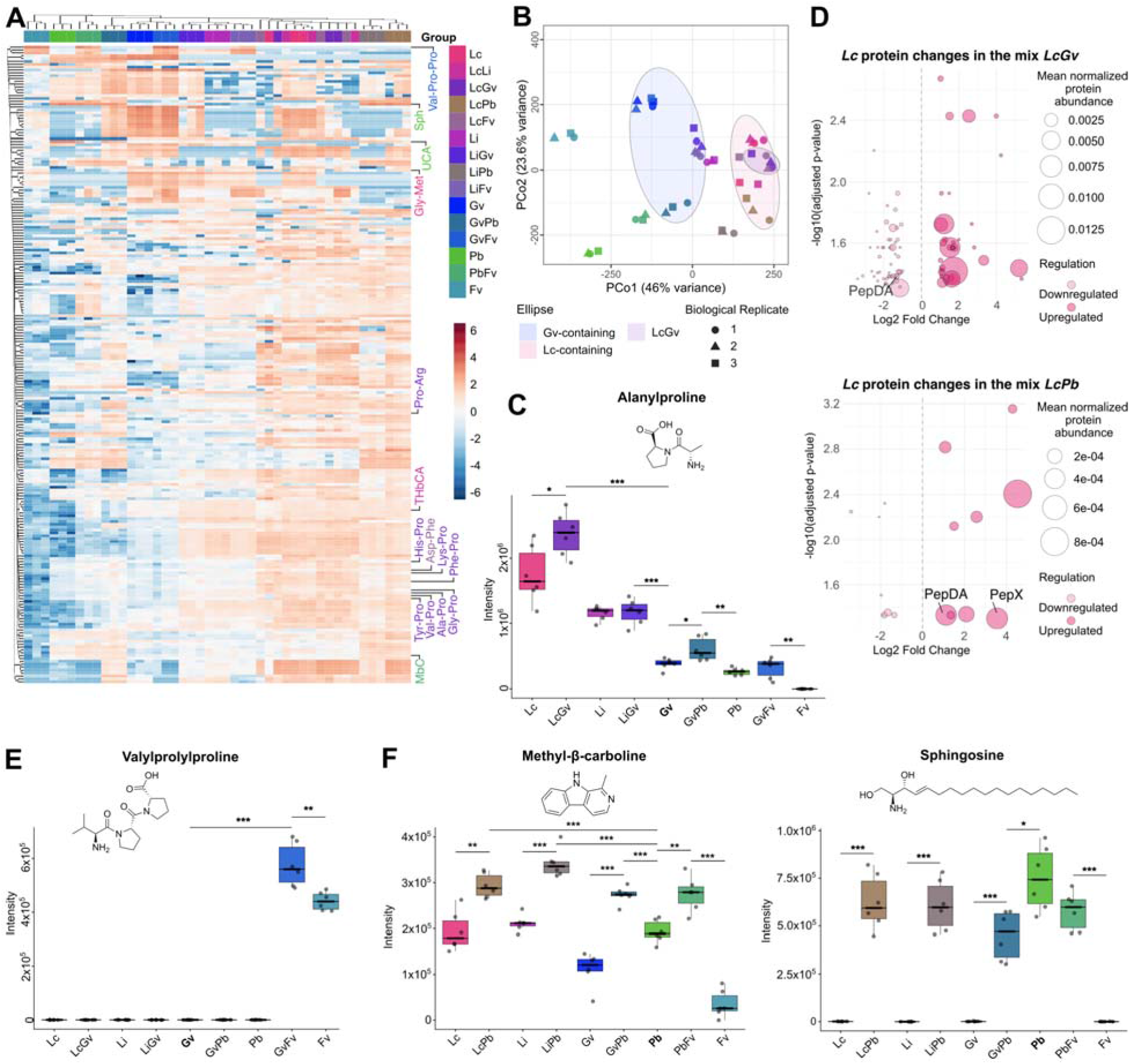
Untargeted metabolomics reveal dipeptides and other metabolites changed in co-cultures. A) Unsupervised hierarchical clustered heatmap of co-culture conditions and all identified metabolite features. Metabolite labels are coloured by the condition where each showed highest intensity. B) Principal coordinates analysis of bacterial metabolite profiles. Points are coloured by experimental group and shaped by biological replicate. Ellipses indicate regions for *Lc*⍰containing, *Gv*⍰containing, and *LcGv* groups. C, E and F) Bar plots show intensity distributions for the indicated metabolite across relevant conditions. Individual biological (n=3) and technical (x2) replicates are shown as black points. Boxes are colored by condition; whiskers extend to the most extreme values within 1.5 x the interquartile range (IQR). The median is marked by a horizontal crossbar. Chemical structures of metabolites are displayed on top of each plot. Statistical comparisons (Welch’s *t*⍰test, *FDR*⍰adjusted) are indicated where applicable (\**p*<0.05, \*\**p*<0.01, \*\*\**p*<0.001). D) Bubble plots showing *Lc* protein changes in the co-cultures *LcGv* and *LcPb*. Downregulated proteins are on the left side in light colours and upregulated proteins on the right side. Circle size indicates the mean normalized protein abundance. Relevant proteins are labelled, (FC > 1) adjusted *p-value* (< 0.05) (Welch’s t-test followed by a Benjamin-Hochberg multiple test correction). Strain abbreviations and colors: *Lc = Lactobacillus crispatus*. Full metabolite names for abbreviations used in the figure ar provided in Extended Data Table S3.

### Interaction-derived metabolites drive divergent Inhibitory outcomes in the vaginal microbiota and multidrug-resistant pathogens

To assess the functional impact of these metabolites on vaginal microbes and broader multidrug-resistant pathogens, we supplemented culture medium with each compound and monitored growth of *Lc*, *Gv*, *Gp*, *Pb*, *Candida albicans* (*Ca*)*, Escherichia coli* (*Ec*), and *Staphylococcus aureus* (*Sa*). Growth curve analysis revealed distinct, species-specific profiles. By calculating the area under the growth curve (AUC), we found that alanylproline (*Gp*, *p* = 0.028, Cohen’s d [*d*] = 3.374), glycylmethionine (*Gp*, *p* = 0.008, *d* = 5.835; *Gv*, *p* = 0.002, *d* = 5.570), and histidylproline (*Gp*, *p* = 0.016, *d* = 4.805; *Gv*, *p* = 0.001, *d* = 6.836) significantly promoted the growth of both *Gardnerella* species. Additionally, histidylproline (*Pb*, *p* = 9×10^-5^, *d* = 27.16), urocanic acid (*Gv*, *p* = 0.008, *d* = 4.858; *Pb*, *p* = 0.001, *d* = 6.781), and aspartylphenylalanine (*Gv*, *p* = 0.002, *d* = 7.201; *Pb*, *p* = 5×10^-4^, *d* = 13.24) increased proliferation of *Gv* and *Pb* (Fig. S7a, b). In contrast, tetrahydro-*β*-carboline-carboxylic acid (1 mM) significantly inhibited *Lc* growth (*p* = 6×10^-4^, *d* = -7.999; Fig. S7b). Harmane (1 mM), a representative methyl-*β*-carboline compound, significantly inhibited the growth of *Gv* (*p* = 5×10^-8^, *d* = -7.069), *Gp* (*p* = 9×10^-4^, *d* = -1.940), *Pb* (*p* = 1×10^-5^, *d* = -3.761), *Ca* (*p* = 2×10^-8^, *d* = -10.28), *Ec* (*p* = 9×10^-7^, *d* = -8.533), and *Sa* (*p* = 0.002, *d* = -2.485), with a weaker effect on *Lc* (*p* = 0.014, *d* = -1.484). It also showed a trend toward reduced biofilm formation in *Gardnerella* (Gp, Cliff’s-δ [*Cδ*] = -0.375; Gv, *Cδ* = -0.458), while significantly preventing it in *Ec* and *Ca* (*p* = 0.005, *Cδ* = -1.000, both). Sphingosine (0.1 mM) significantly inhibited growth of all tested microbes (*p* = 7×10^-14^ - 0.006-, *d* = -16.01 to -2.580) and, except for *Lc*, it also suppressed biofilm formation (*p* < 0.001-0.036, *Cδ* = -1.000 to 0.750; Fig. 7A-D). Together, these findings demonstrate that species-specific metabolic interactions, including growth promotion by the bacteria-derived dipeptides and growth inhibition by methyl-*β*-carboline and sphingosine, contribute to shaping competitive dynamics across microbial communities, from the vaginal microbiota to multidrug-resistant pathogens. They also pinpoint key bioactive molecules that may drive health-dysbiosis transitions or selectively inhibit pathogens, representing potential lead structures for developing novel, selective antimicrobials.

**Figure 7.**
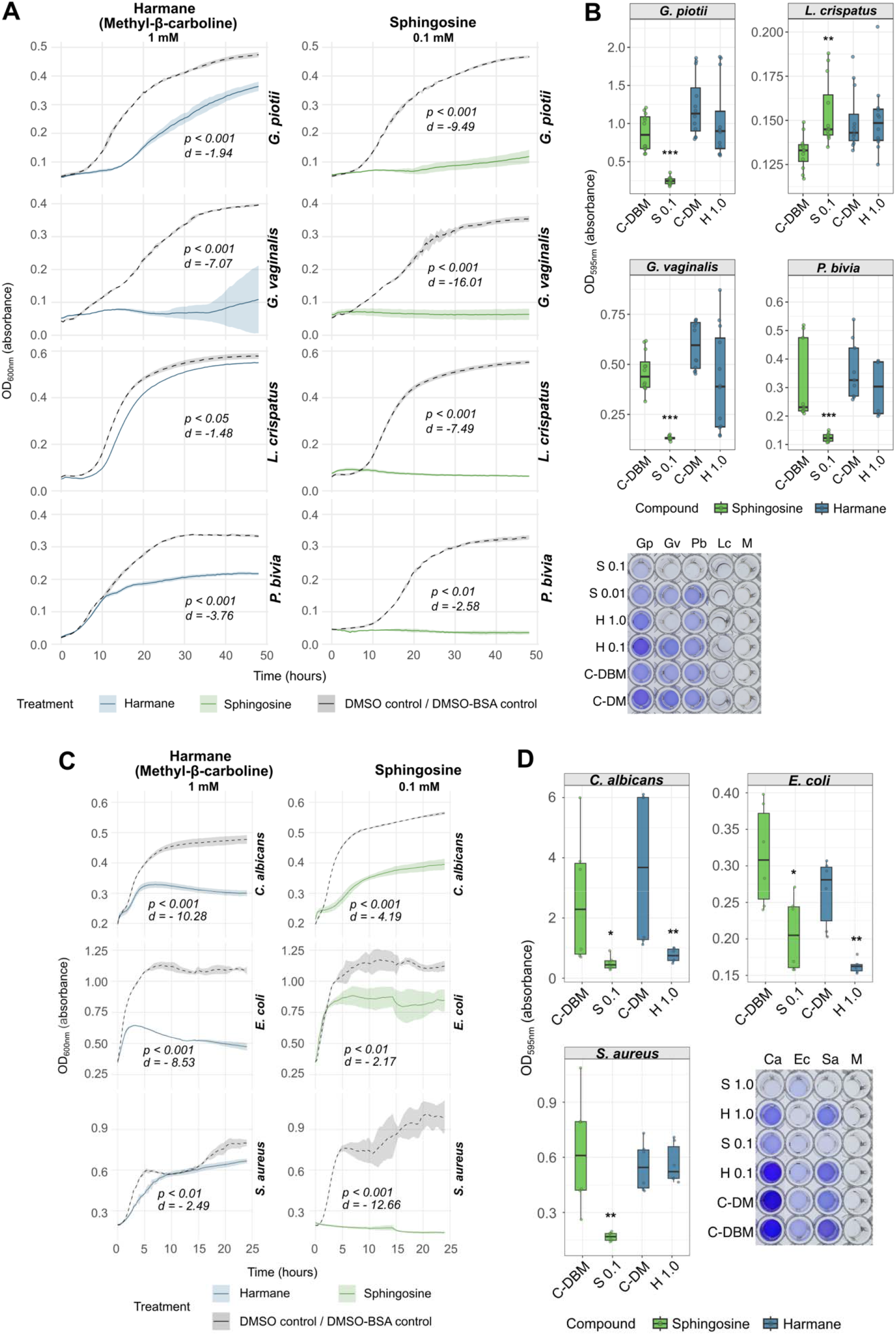
Integrated metabolomic and functional screening identifies metabolites with inhibitory activity against BV-related bacteria. A) Growth curves (Optical density [OD]_600_nm) of vaginal-related bacteria cultured with 1mM of harmane or 0.1 mM of sphingosine. A single curve is shown per condition/strain, representing the mean of three biological replicates, each assayed thrice (except for Pb with sphingosine, n = 2). Alpha-channel-coloured areas correspond to the standard deviation. Statistical significance between treatment and control AUC groups is shown using unpaired two-tailed Student’s t-tests, with effect sizes reported as Cohen’s d. B) Biofilm formation (OD_595_nm) for relevant bacteria treated with sphingosine (0.1 mM) or harmane (1.0 mM) and controls. Points indicate individual replicates. Boxes are colored by condition and whiskers extend to the most extreme values within 1.5 x the interquartile range (IQR). The median is marked by a horizontal crossbar Significance vs. control: \**p* < 0.05, \*\**p* < 0.01, \*\*\**p* < 0.001 (Wilcoxon test, BH correction). Representative image of plate wells for each condition and the corresponding controls are shown. C) Growth curves of multidrug resistant pathogens using same parameters as A. D) Biofilm formation of multidrug resistant pathogens using same parameters as B. Representative image of plate wells for each condition and the corresponding controls are shown. Abbreviations: *Lc = Lactobacillus crispatus, Li = Lactobacillus iners, Gv = Gardnerella vaginalis, Fv = Fannyhessea vaginae, Pb = Prevotella bivia, Ca = Candida albicans, Ec = Escherichia coli, Sa = Staphylococcus aureus and M = culture medium*. C-DM = medium with DMEM, C-DBM = medium with DMEM+BSA, S 0.1 = Sphingosine 0.1 mM and H 1.0 = Harmane 1.0 mM.

## Discussion

In this study, we established a functional genomic resource for the vaginal microbiome by generating high-quality genomes from clinical isolates of *Gardnerella* and *L. iners*. By integrating phenotypic and multi-omics analyses, we identified interspecies genetic variation, particularly in accessory genomic regions. Our findings revealed a conserved *T2SS/Tad* system in *Gardnerella,* and validated its *BV* association with metagenomic cohort data. We also demonstrated that microbiota-derived metabolites inhibit the growth of *BV*-associated bacteria and multidrug-resistant pathogens. Together, these findings underscore the importance of vaginal microbiome research and its translational potential for future diagnosis and therapy.

Previous genomic comparisons between *Gv* strains have revealed significant differences in metabolic and virulence potential [30]. Moreover, pangenome analyses have demonstrated that lateral gene transfer is common within the species [31], and identified heightened metronidazole resistance in certain closely related genospecies [32], linking an evolutionary genomic feature to a clinically relevant resistance phenotype. Our comprehensive pangenomic resource expands the repertoire of genomic-phenotypic associations, revealing a link between biofilm capacity and species divergence, as well as a large cloud-pangenome partition (41% of the total) associated with horizontal transfer, prophage or plasmid genes.

The prevalence of *Gp* in *BV* remains poorly characterized, largely because its recent separation from *Gv* subtypes has limited large-scale epidemiological studies. Nevertheless, higher *Gardnerella* diversity has been associated with increased risk of *BV* [33], suggesting that individual species such as *Gp* may play distinct roles in disease etiology. Moreover, significant associations between Nugent category and the relative abundances of *Gv*, *G. swidsinskii*, and *Gp* have been reported in a co-occurrence study [34], highlighting the importance of accounting for species-level ecological patterns. Notably, our data provide novel insights by defining the prevalence and abundance of both *Gp* and *Gv* in the metagenomic cohort data, together with their biofilm phenotypes, suggesting a mechanism for *BV* in settings where *Gp* co-occurs with other species.

Prior research using dual-species biofilms has revealed the species-specific spatial distribution of bacterial community members [35]. This complexity is underscored by observations that monospecies *Gv* biofilms exhibit reduced transcripts levels for glucose and carbon metabolism [36], while polymicrobial biofilms comprising *Gv, Fv*, and *Pb* show similar downregulation of metabolism-associated genes [37]. Our study employed proteomics in dual-species co-cultures and revealed that regulation depended on both the carbohydrate-scavenging capacity of each species and the identity of the co-culture partner, providing a detailed characterization of partner-dependent metabolic regulation.

A key finding of this study is the discovery of a conserved and potentially functional *T2SS/Tad* operon across all *Gardnerella* strains examined. To our knowledge, this represents the first identification of a *T2SS/Tad* complex in this genus and supports previous reports demonstrating pili on the *Gv* surface [38],[39]. Furthermore, our integration of genomics and AI-predicted structural modeling provides an initial framework implicating this apparatus in DNA uptake or colonization [40],[41],[42]. Notably, specific alleles of this system, mostly in high-biofilm *Gp* strains, suggest it is a potential determinant of *BV* pathogenesis. As surface-exposed and species-conserved machinery required for fitness, the *T2SS/Tad* may offer an attractive target for therapeutic strategies aimed at disrupting biofilm integrity while avoiding off-target effects of broad-spectrum antibiotics. Moreover, the observed *BV*-specific enrichment of the *T2SS/Tad* gene cluster in our cohort analysis supports its potential as a targeted diagnostic tool, particularly in rural settings where sequencing equipment may not be available [43],[44],[45].

Previous studies have examined surface-exposed proteins in *Gv* using cell-shaving proteomics [46]. Quantitative proteomics has also revealed that a low-pH environment triggers upregulation of efflux transporters and virulence factors while downregulating biofilm-associated genes [47]. Additionally, proteomics of cervicovaginal lavages revealed that *Li* and *Lc* proteins involved in DNA protection and glycolysis are significantly reduced in dysbiosis [48]. To our knowledge, our study is the first to perform high-throughput proteomics on a complete all-pairwise co-culture of five key vaginal commensals. In our models, *Gv* downregulated key proteins involved in biofilm dispersal (SufS) [49] and virulence (NanH) while upregulating diverse sugar and nucleotide transporters and lactose utilization when co-cultured with *Lc*, suggesting a metabolic trade-off in which *Gv* prioritizes nutrient scavenging over canonical virulence. In response, both *Lc* and *Li* reduced DNA protection (YbaB/EbfC), while *Lc* upregulated DNA repair/recombination (AddA, RecR, RecA, RecJ) and *Li* increased protein translation (RpsQ, RplD, RplM, RpsJ). This divergence may underlie their differing abilities to withstand *Gv* competition and maintain dominance in healthy versus dysbiotic communities.

Few studies have investigated high-throughput metabolomics of vaginal bacterial interactions. An *in silico* metabolic network of the *Gardnerella* pangenome revealed that genetic relatedness does not predict functional relatedness and identified four core genes essential for metabolic function [50]. A study of microbial diversity and the metabolome associated with *BV* revealed reduced lactate and elevated amines and short-chain fatty acids [51]. Our analysis identified specific compounds produced through interspecies interactions, particularly proline-based di-and tripeptides, alongside regulation of *Lc* dipeptidases (PepDA [52] and PepX [53]) likely involved in their production. Furthermore, methyl-β-carboline-derived metabolites and sphingosine demonstrated selective growth and biofilm inhibition against both vaginal bacteria and multidrug-resistant pathogens. These findings are supported by reports demonstrating antibacterial activity for β-carbolines [54] and sphingosine [55], suggesting that the vaginal microbiome constitutes an untapped reservoir for novel antimicrobial scaffolds.

This study has several limitations. First, the limited number of isolates per category precluded statistical analysis of potential associations between biofilm-forming capacity and the available clinical metadata [56],[57],[58]. Second, although AI multimer predictions provided structural insights into the *T2SS/Tad* complex, the moderate interface confidence scores warrant cautious interpretation, underscoring the need for high-resolution experimental validation of the proposed quaternary architecture. Third, our *in vitro* interaction and inhibition models require further validation in more complex systems, such as polymicrobial communities or host-microbe co-cultures. Finally, future work should investigate how the bacterial genes, proteins, and metabolites identified here modulate host epithelial and immune responses, thereby bridging microbial ecology with host pathophysiology.

In conclusion, this work employed integrated multi-omics and functional analyses, including 16S rRNA gene sequencing, genomics, proteomics, and metabolomics, to investigate key vaginal microbiome species and their interactions. It establishes a model in which divergence in *Gardnerella* biofilm capacity is genetically encoded within an accessory genome, functionally deployed through tailored proteomic responses to interspecies cues, and further modulated by an interaction-specific metabolic landscape. Together, these findings provide a foundation for identifying bioactive molecules in the vaginal microbiome and for developing diagnostics and targeted therapeutics to restore and maintain vaginal health.

## Methods

### Microbial strains

For this project, we used both clinical isolates and reference strains. NYC III medium supplemented with 10% FBS was used to culture all bacteria on both agar plates and in broth under anaerobic conditions at 37°C. Clinical isolates were collected from healthy individuals with information on HPV status, and the vaginal microbiota community type [57],[58]. All microbial clinical stocks were stored in glycerol solution at -80°C prior to usage. A total of 18 different clinical isolates were used: *Gardnerella vaginalis* strains 175, 144-3, 144-4, 95, and 88; *Gardnerella piotii* strains 133, 133-1, 133-2, 142-2, and 142-3, *Gardnerella pickettii* strain 221, *Lactobacillus iners* strains H4, H12, H30An, H30Ae, H32, H78An, and H78Ae. Reference strains were obtained from Culture Collection University of Gothenburg (CCUG) and American Type Culture Collection (ATCC). The following 12 reference strains were used during this study: *G. vaginalis* CCUG 44012, CCUG 44069, CCUG 44089, and CCUG 44114; *L. iners* CCUG 44065, CCUG 44108, CCUG 46933, and ATCC 55195; *Lactobacillus crispatus* CCUG 44500, *Fannyhessea vaginae* CCUG 44116, *Prevotella bivia* CCUG 9557T. Reference strains were derived from healthy individuals, individuals with BV, or individuals with other health issues (Supplementary Tables 1a). In addition, *C. albicans* (CCUG 44135), and antibiotic-resistant pathogens *E. coli* (ST131 BE05A 16, resistance to Ampicillin, Cefotaxime, Ciprofloxacin, nalidixic acid, and Cefpodoxime) and *S. aureus* (ATCC 29213, resistance to Ciprofloxacin, Clindamycin, Gentamicin, and Neomycin) were used.

### Preparation of bacterial cultures and crystal violet assay

NYC III medium broth and agar plates were deoxygenated inside the anaerobic chamber for at least 24 h before use. Biofilm formation was assessed using a crystal violet assay. Briefly, *Gardnerella* and *L. iners* were cultured anaerobically on agar and subsequently in NYCIII broth. Following 24-48 h growth, bacterial cultures were normalized to OD_600_ = 1.0 with fresh medium. Aliquots (200 µL) of standardized suspensions were inoculated in triplicate into a 96-well plate and incubated anaerobically for 24–48 h at 37°C. Adherent biofilms were washed with PBS, stained with 0.1% (w/v) crystal violet, solubilized with 30% acetic acid, and quantified by measuring OD_595_ (SpectraMax i3x, Molecular Devices. San Jose, CA, USA) in 96-well plates (100 µL). Data were normalized by subtracting the average absorbance of NYCIII medium-only controls, and data analyses were performed using *dplyr* (v1.1.4) and *tidyr* (v1.3.1) packages in R (v4.4.2).

### DNA extraction, quality control and whole genome long-read sequencing

Strains were selected for DNA extraction and long-read sequencing based on the relatively high or low biofilm formation compared to other strains of the same species. Reference strains used for proteomics analysis were selected based on their relevance reported in literature [3, 35, 37]. DNA extraction was carried out using the Quick-DNA™ High Molecular Weight MagBead Kit (Zymo Research. Irvine, CA, USA) following the provided protocol. Quality control of the extracted DNA included measuring DNA purity, concentration and fragmentation using NanoDrop™ 2000 Spectrophotometer (Thermo Fisher Scientific. Waltham, MA, USA), Qubit 4 Fluorometer (Invitrogen. Carlsbad, CA, USA), and 4200 TapeStation System (Agilent Technologies. Santa Clara, CA, USA), respectively. Extracted DNA was prepared for long-read sequencing with the Rapid Sequencing Kit or the Rapid Barcoding Kit 24 V14 (Oxford Nanopore Technologies. Oxford, UK), following the respective protocols. Nanopore sequencing was carried out using MinION R10.4.1 and Flongle R9.4.1 Flow Cells (Oxford Nanopore Technologies. Oxford, UK). Software used during the sequencing runs were MinKNOW (v22.05.05-24.02.6), MinKNOW core (v5.1.0-5.9.7), Bream (v7.1.3-7.9.5), and Configuration (v5.1.5-5.9.13). Basecalling was performed using Guppy (v6.1.5-6.4.6) or Dorado (v7.2.13-7.3.9), set to super-accuracy or high-accuracy basecalling mode.

### Downstream analysis of sequencing data

Using conda (v24.1.2) and chopper [59] (v0.7.0-0.8.0), .fastq files obtained after sequencing were filtered and trimmed for a minimum Phred average quality score of 12. Flye [60] (v2.9.3-2.9.4) was used for *de novo* assembly, and proovframe [61] (v0.9.7) and DIAMOND [62] (v2.1.9.163) for frame-shift correction. *Gardnerella vaginalis* FDAARGOS_568 genome assembly ASM381276v1, *Lactobacillus iners* C0322A1 genome assembly ASM1105877v1, *Fannyhessea vaginae* NCTC13935 genome assembly 58174_B01, *Lactobacillus crispatus* FDAARGOS_743 genome assembly ASM973027v1 from the National Center for Biotechnology Information were used as reference genomes for aligning with proovframe. All assemblies used for post-processing were single-chromosome circular contigs. With the exception of *G. vaginalis* 142-3, only assemblies with at least 45x coverage were included in downstream analysis [63]. Genome completeness was assessed using CheckM [64] (v1.2.3–1.2.4), ranging from at least 96.41% to 100%. Genome annotation was performed using the Prokaryotic Genome Annotation Pipeline Annotation Software PGAP [65] (revision 2024-07-18.build7555 and 2025-05-06.build7983; and revision 6.11). The annotated amino acid fasta files were used for pangenome analysis with Partitioned PanGenome Graph Of Linked Neighbors PPanGGOLiN[66] (v2.3.0) using default settings. Additional reference annotated amino acid fasta files for pangenome analysis were retrieved from the National Center of Biotechnology Information (NCBI), focusing on genomes with a minimum completeness of 96.41%. Pangenome graphs were generated using Gephi [67] (v0.11.2). Gene maps were generated using *gggenes* (v0.5.1) under *ggplot2* package (v4.0.0). Gene identity was obtained using *stringr* (v1.5.2), *pwalign* (v1.2.0), and *Biostrings* (v2.74.1) packages. Module prediction was performed using the panModule [68] workflow included in PPanGGOLiN and run with default parameters.

### Prophage identification and quality assessment

Prophage regions were identified in assembled *Gardnerella* genomes using geNomad v1.12.0 and extracted from corresponding contigs. Five candidate regions were retained: two from Gv88 and one each from Gv142-2, Gv144-3, and Gv44089. Completeness and contamination were assessed with CheckV, and predictions were compared with PHASTEST where available. Two references were included: *Gardnerella* phage vB_Gva_AB1 (GenBank MW387018.1) and a pGamma prophage region reconstructed from the chromosome of pGamma-positive isolate DNF00622A, the latter treated as a reconstructed reference prophage. Candidate and reference sequences were annotated with Pharokka v1.10.1. CDSs, phage-associated functions, and terminase large-subunit (TerL) proteins were retrieved, and similarity to INPHARED phages was assessed via Mash. Functional genome maps were generated from Pharokka annotations using pyCirclize v1.10.1. CDSs were displayed by transcriptional orientation and colored by functional category, with representative phage-associated genes highlighted where present. Local GC content and GC skew were calculated with a 500-bp sliding window (100-bp step) and displayed as internal tracks. TerL amino-acid sequences were extracted from each candidate and reference sequence. Representative homologues were identified by BLASTp against NCBI ClusteredNR, selecting full-length or near-full-length sequences. The final dataset comprised 19 TerL sequences (five candidate prophages, two references, 12 public homologues). Sequences were aligned with MAFFT v7.526 (L-INS-i), trimmed with trimAl v1.5.rev1 (-automated1), and analyzed by maximum-likelihood inference in IQ-TREE v3.1.3. ModelFinder selected Q.PFAM+G4 by BIC. Branch support was assessed with 1,000 ultrafast bootstrap and 1,000 SH-aLRT replicates, and the topology was displayed as a topology-only cladogram with node support shown only when both values were ≥ 80%. As all candidate prophage sequences were computationally extracted from bacterial chromosomes, they were treated as predicted prophage regions rather than experimentally validated complete phage genomes. Sequence orientation depended on host contig orientation, so reverse-orientation alignments were not interpreted as biological inversions, and circular layouts were used solely for visualization, not as evidence of circular genome topology. Analyses used geNomad v1.12.0, Pharokka v1.10.1, VClust v1.3.1, BLAST+ v2.17.0, MUMmer v4.0.1, clinker v0.0.32, MAFFT v7.526, trimAl v1.5.rev1, IQ-TREE v3.1.3, and pyCirclize v1.10.1. Custom Python scripts summarized pairwise ANI/AF values and generated comparative heatmaps and functional genome visualizations.

### Pairwise bacterial co-culture experiment

Five key vaginal bacterial species (*L. crispatus* CCUG 44500, *L. iners* CCUG 44108, *G. vaginalis* CCUG 44012, *F. vaginae* CCUG 44116, and *P. bivia* CCUG 9557T) were initially streaked separately on NYCIII agar and incubated anaerobically for 48 h. For co-culture experiments, strains were normalized to OD_600_ = 1.0 and combined 1:1 in dual cultures in 2 mL tubes to perform all pairwise combinations of the five species, including monoculture controls. Following centrifugation, pellets were resuspended in fresh NYCIII broth to a final volume of 4 mL per replicate. After 24 h of anaerobic incubation, cultures were pelleted via centrifugation. Supernatants were filtered (0.2 µm), mixed with 4 volumes of ice-cold methanol, and stored at - 80°C until the metabolomic analysis. Cell pellets were washed three times with PBS and stored dry at -80°C for subsequent proteomic and 16S rRNA gene sequencing analysis.

### Proteomics sample preparation and digestion

For proteomics analysis, 70% of the pellets from the co-culture experiment were collected and submitted to Nanoxis Consulting AB, Gothenburg, Sweden. Bacterial pellets were lysed by bead-beating for 5 min at 30 Hz in PBS containing 2% SDS using acid-washed glass beads (TissueLyser II, QIAGEN. Hilden, Germany). Supernatants were collected, and protein concentration was determined via BCA assay (Thermo Fisher Scientific. Waltham, MA, USA). Proteins were digested using either a modified filter-aided sample preparation (FASP) protocol on 30 kDa MWCO Nanosep 30k Omega filters (Merck) or an adapted SP3 method through precipitation on Sera-Mag™ SpeedBeads (Carboxylate-Modified, Cytiva). Protein samples were reduced with 10 mM DTT at 60°C for 30 min and alkylated using either 10 mM methyl methanethiosulfonate or 40-50 mM iodoacetamide for 30 min at room temperature. Proteins on FASP filters were washed with 8M urea and digested in 1% sodium deoxycholate in 50 mM triethylammonium bicarbonate (TEAB). Proteins precipitated and washed on SP3 beads with 70% ethanol and resuspended in 50mM TEAB for digestion. LysC/Trypsin was added (1:50 enzyme to protein ratio, Promega) and incubated o/n at 37°C. Additional trypsin (1:100, Thermo Fisher Scientific) was added and incubated for another two hours. Peptides were collected and cleaned up using HiPPR Detergent Removal Kit and Pierce Peptide Desalting Spin columns (both Thermo Fisher Scientific) according to the manufacturer instructions. LCMS analyses were performed on an Orbitrap Tribrid (Lumos/Eclipse) or on an Orbitrap Exploris mass spectrometer, which were interfaced with an Easy-nLC1200 liquid chromatography system (all Thermo Fisher Scientific). Peptides were trapped on an Acclaim Pepmap 100 C18 trap column (100 μm x 2 cm, particle size 5 μm, Thermo Fischer Scientific) and separated on an in-house packed analytical column (75 μm x 35 cm, particle size 3 μm, Reprosil-Pur C18, Dr. Maisch) using a gradient from 5% to 80% acetonitrile in 0.2% formic acid over 90 min at a flow of 300 nL/min. The instruments operated in data-dependent mode where the precursor ion mass spectra were acquired at a resolution of 120 000 and m/z ranging from 375 to 1500. Precursors with charge states 2 to 6 were selected for fragmentation using 30% (HCD) or 35% (CID) collision energy. The isolation window was set between 0.7 to 1.4 Da and dynamic exclusion to 30 s to 60 s and 10 ppm. MS2 spectra were recorded at a resolution of 30 000 with maximum injection time set to 110 ms by Orbitrap or by IonTrap at rapid scan rate with a maximum injection time of 50 ms.

### Proteomics raw data analysis

Acquired raw data files were analyzed using Proteome Discoverer v3.0 (Thermo Fisher Scientific) with MS Amanda v2.0 search engine against a custom-designed database with 10,028 protein sequences. A maximum of two missed cleavage sites were allowed for full tryptic digestion, while setting the precursor and the fragment ion mass tolerance to 10 ppm and 0.6 Da, respectively. Tryptic peptides were accepted with 1 missed cleavage; methionine oxidation was set as a variable and cysteine carboxymethylation was set as fixed modifications. For PSM validation, Fixed Value PSM Validator or Percolator (1% FDR) were chosen. *In house* amino acid annotated reference strains sequences were provided for using as database. Sequences were obtained as described above, except for *L. crispatus* VPI 3199 (ATCC 33820) and *P. bivia* CCUG 9557T (ATCC 29303), which were retrieved from NCBI database (Genome assemblies ASM1898723v1, ASM4042808v1).

### Proteome abundance bioinformatic analysis

Protein metadata and abundance values for the control and mixed conditions were extracted, and only entries corresponding to the target species per analysis were retained. Potential contaminants and proteins with fewer than two unique peptides were removed. Additionally, proteins that were identified but not quantified (zero-abundance proteins) were discarded. Missing abundance values were imputed with half the minimum abundance observed for each protein within species. After filtering out proteins from irrelevant species, protein abundances for each sample were normalized by total protein abundance. Differential expression between co-culture samples and monoculture controls was evaluated using a two-sided Student’s *t*-test with unequal variance for mixed samples versus each control (representing individual organisms). An adjusted *p-value* cutoff of 0.05 with Benjamini-Hochberg for multiple testing was applied to identify significantly altered proteins in each comparison. Strain-unique proteins were defined as protein products inferred from two or more peptide fragments uniquely assigned to a single bacterial strain locus tag. We parsed the protein product lists for each relevant strain under co-culture conditions and identified unique occurrences per strain. For homologous product names within and across strains, the locus_tag differentiates the products. An additional manual assessment at the peptide sequence level was done for random selected homologous products within and between strains. Aside from finding identical peptides with different post-translational modifications within individual strains, no identical peptides were detected between strains. Data analysis was performed using packages *dplyr* (v1.1.4), *tidyr* (v1.3.1), *stringr* (v1.5.2) and *tibble* (v3.3.0) under R (v4.4.2). Additional packages used for visualization were *scales* (v1.4.0) (overall protein regulation), *ggvenn* (v0.1.19) (Venn diagrams), *ggrepel* (v0.9.6) (bubble plots), *ggplot2* (v4.0.0).

### Proteomic shift index calculation

To capture global proteome remodeling across conditions, we calculated a Proteomic Shift Index (PSI) for each treatment. For a given condition, all proteins with a raw *p* < 0.05 (Welch’s t-test) were classified as up- or downregulated based on their log₂ fold change relative to the control; no fold-change cutoff was applied. The PSI was calculated separately for each direction as:

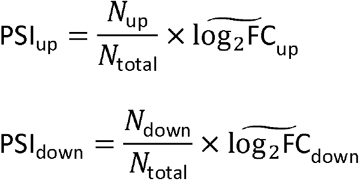

where *N*_up_ and *N*_down_ are the numbers of significantly up- and downregulated proteins, *N*_total_ is the total number of proteins quantified in that condition, and 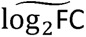 represents the median log₂ fold change of the respective subsets. The median was chosen over the mean to reduce the influence of outlier proteins with extreme fold changes, thereby providing a more robust estimate of the central tendency of each directional shift. PSI_up yields a positive score reflecting the protein fraction-weighted magnitude of upregulation, while PSI_down yields a negative score reflecting the protein fraction-weighted magnitude of downregulation, allowing independent assessment of each directional shift. For the *FvPb* coculture condition, where lower proteome coverage limited statistical power, the significance threshold was relaxed to raw *p* < 0.1 (Welch’s t-test).

### Protein enrichment and network analysis

STRING [69] web server version 12.5 was used to perform functional enrichment analysis of Gardnerella vaginalis CCUG 44012. The complete set of amino acid sequences of this strain was first uploaded to STRING to generate a custom organism entry. For each of the up- and downregulated proteins, a signed rank score was calculated from the log₂ fold-change (log₂FC), and raw *p-value* as follows:

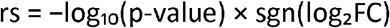

where sgn is the sign function. The resulting locus tag-signed rank score list was then submitted to STRING using the “Proteins with Values/Ranks” input mode for functional enrichment analysis. Default parameters were used, including a false discovery rate (FDR) stringency of 1% (high confidence). For network analysis, the “Multiple proteins” input mode was used with the set of significantly changed proteins, defined as those with |log₂FC| > 1.0 and Benjamini–Hochberg adjusted *p* < 0.05.

### Mapping of differentially expressed proteins in the pangenome

The list of significantly differentially expressed protein products in *G. vaginalis* CCUG 44012 was parsed against the gene list of pangenome modules present in *G. vaginalis* CCUG 44012 and only unique proteins associated with single11copy genes were retained based on the *locus tag* matched.

### AI prediction of *T2SS/Tad* complex

Structural models for core type II secretion Tad pilus complex system (*T2SS/Tad*) and components were predicted using AlphaFold3 [70]. Amino acid sequences of homologous TadA family conjugal transfer-associated ATPase and type II secretion system F family protein (s) served as input queries. Assembly of the *T2SS/Tad* complex was based on the canonical architectural framework described by Pelicic [27], assuming a hexameric ATPase ring. For the platform subcomplex, multiple stoichiometrically plausible configurations of the T2SS F family protein were tested. The final model was selected based on its superior AlphaFold3 confidence metrics (ipTM and pTM scores) among all tested configurations. Transmembrane domains were predicted using DeepTMHMM [71] (v1.0.44) via the Technical University of Denmark Bioinformatic Server (https://dtu.biolib.com/DeepTMHMM/). Functional domains were predicted using InterPro [72] 107.0 (https://www.ebi.ac.uk/interpro/) For structural analysis and figure generation, protein models were visualized, rendered, and aligned using PyMOL Molecular Graphics System (v3.1.0), Schrödinger, LLC.

### 16S rDNA gene sequencing and data analysis

The remaining 30% of the pellets from the co-culture experiment were collected and resuspended in PBS for DNA extraction using the Quick-DNA^TM^ Miniprep Kit (Zymo Research. Irvine, CA, USA) following the standard protocol for bacteria cell wall lysis. DNA samples were submitted to Biomarker Technologies (BMKGENE. Münster, Germany) for 16S rDNA sequencing. Briefly, Illumina Novaseq platform was used for generating paired-end reads of the targeted 16s v3+v4_b regions (F:ACTCCTACGGGAGGCAGCA; R:GGACTACHVGGGTWTCTAAT). Raw reads were quality-trimmed using Trimmomatic (v0.33; sliding window: 50bp, Q20) and adapters/primer sequences were removed with Cutadapt (v1.9.1; max error: 0.2). Paired-end reads were merged (USEARCH v. 10; min overlap: 10bp, min identity: 90%), and chimeras were filtered using UCHIME (v8.1). Sequence variants were inferred using the DADA2 pipeline in QIIME2 (v2020.6). In parallel, OTUs were clustered at 97% similarity with USEARCH (v10.0). Both ASVs and OTUs were filtered at a 0.005% minimum relative abundance threshold across all samples. Taxonomic annotation was performed using a combined approach. Sequences were first compared via BLAST against the SILVA (v138), UNITE (v8.0), and Greengenes (v13.5) databases using classify-consensus-blast in QIIME2 (min identity: 90%, min coverage: 90%). Unassigned sequences were subsequently classified with a naive-Bayes classifier (classify-sklearn; confidence threshold: 0.7) trained on the same reference databases. Obtained relative abundance was filtered and normalized based on 16S rDNA copy number per bacterial species based on The ribosomal RNA Database rrnDB[73] (https://rrndb.umms.med.umich.edu/genomes/?order_by=-dsorgname) using the formula:

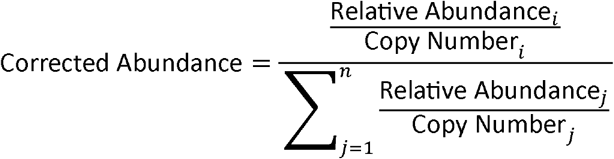

Where:

- *i* = the specific taxon calculating for
- *j* = all taxa in the sample
- *n* = total number of taxa

Although accounting for 16S rDNA copy number variation is challenging in metagenomic surveys of unknown communities [74], this adjustment provides a more accurate estimate of relative bacterial abundance in our setting, where the sequence-annotated bacterial strains are well characterized.

### T2SS/Tad abundance data analysis from clinical cohorts

A total of 125 metagenomic samples were included in this study, comprising 43 samples from patients with bacterial vaginosis (*BV*) and 82 samples from healthy controls. Following downstream filtering for *T2SS/Tad* module completeness, 93 samples (39 *BV* and 54 healthy controls) were retained for the final quantitative analysis. Raw paired-end metagenomic reads were quality-filtered to remove low-quality bases and adapters. Host contamination was removed by aligning reads to the human reference genome (T2T-CHM13v2.0) using Bowtie2 (v2.5.4), and only unmapped reads were retained for downstream analysis. A custom protein database was constructed using five core genes of the putative *T2SS/Tad* module (TadA family conjugal transfer-associated ATPase, type II secretion system F family protein, type II secretion system F family protein, TadE/TadG family type IV pilus assembly protein, Rv3654c family TadE-like protein). Housekeeping genes were excluded to ensure specificity. Host-depleted reads were concatenated and aligned against this database using DIAMOND (v2.0.4) in blastx mode with the following parameters: -evalue 1e-5, -id 50, -query-cover 70, and -max-target-seqs 1. To ensure functional relevance, samples with ≥ 60% of core module genes detected were retained. *T2SS* abundance was quantified as Reads Per Million (RPM), calculated as the number of unique reads mapped to the core genes divided by total clean microbial reads per sample and multiplied by 10⁶. Statistical analyses were performed in R (v 4.4.0). Differences between *BV* and healthy groups were assessed using the Wilcoxon rank-sum test. Data visualization was performed using *ggplot2* and *ggpubr*. A *p*-value < 0.05 was considered statistically significant.

### Metabolomics sample preparation

For LC-MS analysis, 500 μL of methanol-preserved supernatant samples were collected after centrifugation, and 10 μL of every sample was aliquoted into a QC Eppendorf tube. 400 μL of supernatant were transferred into new Eppendorf tube and 20 μL of internal standard mixture was added. The mixture was dried under vacuum in the SpeedVac. The dried pellets were redissolved in 50 μL of 5% MeOH solution. Internal standard mix consist of next isotopically labeled compounds: Tyrosine (5 μg/mL), Phenylalanine (10 μg/mL), and Valine (20 or 30 μg/mL).

### Metabolomics UHPLC–MS/MS analysis

The UHPLC–MS/MS analysis was performed in a Maxis II ETD Q-TOF mass spectrometer (MS, Bruker Daltonics, Germany) using an electrospray ionization (ESI) source with an Elute UHPLC system (Bruker Daltonics, Germany). The separation was performed on an Acquity UPLC HSS T3 column (1.8 μm, 100 × 2.1 mm) from Waters Corporation. Milli-Q water with 0.1% FA was used as mobile phase A, and MS grade methanol with 0.1% FA was used as mobile phase B. The column temperature was kept at 40 °C, and the autosampler temperature was kept at 5 °C. The flow rate was set to 0.2 mL/min with an injection volume of 5 μL. The gradient used was as follows: 0–2 min, 0% B; 2–15 min, 0–100% B; 15–16 min, 100% B; 16–17 min, 100–0% B; 17–21 min, 0% B. The system was controlled using the Compass HyStar software package from Bruker (Bruker Daltonics, Germany). High-resolution mass spectra were acquired in positive and negative mode at a mass range of *m*/*z* 50–1,200. Data acquisition was performed in AutoMSMS mode (data-dependent acquisition, DDA) with a cycle time of 0.5 s and a ramped collision energy from 20 to 50 eV. A solution of sodium formate [10 mM in a mixture of 2-propanol/water (1/1, v/v)] was used for internal calibration at the beginning of each run, in a segment between 0.1 and 0.3 min. The samples were injected to the UPLC-MS system in a randomized order with QC samples injected in the beginning and end of the sample list in both ionization modes, as well as after every eight samples. The standard deviation of the calibration for all injections was below 0.5 ppm.

### Chromatograms and MS data analysis

The chromatograms and mass spectra were processed using the XCMS R package (v4.2.1) for peak detection and retention time correction, focusing on a single peak in both positive and negative ionization modes [75, 76]. After XCMS, Systematical Error Removal using Random Forest (SERRF) normalization was performed using R script (Shiny-SERRF from UCDavis) [77]. Features with intensities > 20,000 ion count, retention time > 1 and < 18 min, and CV% of the QCs < 30 were selected for further statistical analysis. The intensities of the internal standards and the QC samples were plotted against the UPLC-MS/MS sample injection order to evaluate the stability and performance of the experimental set over time. For hypothesis testing, features were first filtered based on fold-change, requiring all co-cultures of interest to differ from the corresponding single-culture control by at least 1.5-fold in the same direction (increase or decrease). Two-tailed *t*-tests assuming unequal variances (Welch’s *t*-test) were then applied to the filtered features.

### Identification of metabolites

Significant features and molecules of interest were primarily annotated by databases (https://www.hmdb.ca, https://mona.fiehnlab.ucdavis.edu/) based on m/z value, high resolution mass and AutoMSMS fragmentation spectra [78]. The SIRIUS platform was used for spectra screening [79–81]. An MS/MS screening script was developed in R script using MetaboAnnotation and CompoundDb packages [81]. The standards were acquired under the same conditions as the sample acquisition method. The experimental *m/z*, retention time and MS/MS were extracted and exported as mgf file for each standard using DataAnalysis. An internal reference database was created and managed with the CompoundDb package in sqlite format, which allows high level of identification confidence. Meanwhile, MassBank (release 2021-03) was also used to screen the MS/MS [81, 82]. We have used a combination of structure elucidation with our *in house* chemical library of authentic standards as well as the SIRIUS software to classify metabolites into four confidence levels [83]. To identify statistically important up- and downregulated features, we utilized our metabolomics identification pipeline including a high-resolution mass screening of public databases, fragmentation spectra comparison with an in-house library, SIRIUS, and manual structure elucidation.

### Growth and biofilm assays with metabolite supplementation

The following compounds, identified as significantly altered in our metabolomic analysis, were obtained from commercial suppliers (Sigma Aldrich. St. Louis, USA) for functional validation: tetrahydro-β-carboline-carboxylic acid, harmane (to represent methyl-β-carboline), both dissolved in DMSO (working concentration 0.5%), and sphingosine dissolved in DMSO (0.5%) and BSA (4mg/mL). Each compound was tested for its effect on bacterial growth and biofilm formation. Microbial cultures were grown in standard medium supplemented with compounds at 0.01 mM, 0.1 mM or 1.0 mM. Growth was monitored for 48 h or 24 h by measuring OD_600_ every 30 min or 20 min in a Tecan Sunrise^TM^ microplate reader (TECAN. Männedorf, Switzerland) using Magellan software (v7.5). Following the growth assay, biofilm formation was immediately quantified from the same cultures using crystal violet assay described above. To quantify bacterial growth, the area under the curve (AUC) was calculated for each well using the trapezoidal rule[84]. Statistical significance between treatment and control groups was assessed using unpaired two-tailed Student’s t-tests, with effect sizes reported as Cohen’s d. Biofilm formation was compared between each treatment and its matched control using a two-tailed Wilcoxon rank-sum test (Mann–Whitney U, normal approximation due to ties). Effect sizes are reported as Cliff’s delta (δ). The Hodges–Lehmann estimator with 95% confidence intervals is reported for median differences. Additionally, glycylproline, alanylproline, glycylmethionine, valylprolylproline, histidylproline, urocanic acid, and aspartylphenylalanine were also tested on bacterial growth and biofilm formation at 1 mM, 5mM or 10 mM. Data processing and statistical analysis were performed in R (v4.4.2) using the tidyverse packages *dplyr* (v1.1.4), *tidyr* (v1.3.1), *stringr* (v1.5.2), *readr* (v2.1.5), and *broom* (v1.0.10).

All data visualization was performed using *ggplot2* (v4.0.0) within the R environment (v4.4.2, “*Pile of Leaves*”) in RStudio (v2024.09.1+394). Figures were exported as scalable vector graphics using the *svglite* package (v2.2.2). Workflow illustration was generated using BioRender (https://BioRender.com). Additional figure edition was performed using Inkscape (v1.4.2).

### Use of Large Language Models (LLMs)

We used the DeepSeek large language model (DeepSeek-V3.2-Exp - DeepSeek-V4.1-Flash-1017) as an assistive tool during coding development and manuscript preparation. Specifically, pseudocode entries describing each analytical step were provided to the model, which generated draft code implementations; these drafts were subsequently reviewed, tested, and refined iteratively by the authors until the resulting scripts reproduced the intended analyses. In addition, the model was used to check the grammar and syntax of the manuscript text. All LLM-generated output was verified by the authors, who take full responsibility for the correctness and integrity of the code and the final manuscript.

## Data availability

All data analysed during this study are included in this published article and its supplementary information / source data files. Genome assemblies and annotations created from this study are available under the BioProject PRJNA1426591 at the National Center for Biotechnology Information. The mass spectrometry proteomics data have been deposited to the ProteomeXchange Consortium via the PRIDE [85] partner repository with the dataset identifier PXD082932. Metagenomic sequencing data were retrieved from publicly available metagenomes in the NCBI Sequence Read Archive (SRA) under accession numbers PRJNA838641, PRJNA208535, PRJNA1398572, and PRJNA707585.

## Code availability

The scripts to reproduce the data analysis in the article are available at https://github.com/FabricioRGKI/Gardnerella-multiomics, while those required to reproduce the prophage analysis are available at https://github.com/YeLiu-ki/Gardnerella_T2SS.

## Ethics declarations

Clinical bacterial isolates were obtained from previously published cohorts under ethical permissions 2017/725-31, 2019-04201, 2021-01847, 2022-04081-02 and 2023-04703-02 from the Stockholm Regional Ethics Committee.

## Acknowledgements

This study was funded by the Swedish Research Council (ID:2025-02904 to J.D., ID: 2020-04707 to D.G., ID:2023-02692 to A.A.S.), Cancerfonden (ID: 23 2916 Pj to J.D., ID: 25 4898 Pj to D.G.), Radiumhemmets Forskningsfonder (ID:251172 to J.D.), Karolinska Institutet (to J.D.) and Magnus Bergvalls Foundation (ID:2023-898 to A.A.S.). Proteomic analysis was performed at the Proteomics Core Facility, Sahlgrenska academy, Gothenburg University, with financial support from SciLifeLab and BioMS. MALDI-ToF identification of strains was done in collaboration with culturomics team at Translational Microbiome Research and Pandemic Preparedness, Karolinska Institutet, Sweden. *E. coli* (ST131 BE05A 16) and *S. aureus* (ATCC 29213) were obtained from Professor Åsa Sjöling’s group, University of Gothenburg, Sweden.

## Authors contribution

F.R.G., V.D., D.G., and J.D. designed the study and analyzed the data. F.R.G., V.D., S.L.K., K.J.v.D. and C.F., performed the experiments. F.R.G., V.D., A.K., R.K., C.L., D.G. and J.D. contributed to experimental design and data interpretation. F.R.G., V.D., H.G., A.M.M., A.V. and L.W.H. performed omics and statistical analysis. Y.L. performed T2SS/Tad cohort and prophage analysis. A.A.S., D.G., and J.D. obtained funding. L.E., A.A.S., D.G., and J.D. supervised the project. F.R.G. and J.D. wrote the manuscript with input from all co-authors. All authors read and approved the final manuscript.

## References

1. The Human Microbiome Project Consortium. Structure, function and diversity of the healthy human microbiome. Nature. 2012;486:207–14. 10.1038/nature11234.

2. Cheng Q, Lv S, Yin N, Wang J. Microbial regulators of physiological and reproductive health in women of reproductive age: their local, proximal and distal regulatory roles. Npj Biofilms Microbiomes. 2025;11:207. 10.1038/s41522-025-00839-y.

3. Ravel J, Gajer P, Abdo Z, Schneider GM, Koenig SSK, McCulle SL, et al. Vaginal microbiome of reproductive-age women. Proc Natl Acad Sci. 2011;108 supplement_1:4680–7. 10.1073/pnas.1002611107.

4. Ravel J, Moreno I, Simón C. Bacterial vaginosis and its association with infertility, endometritis, and pelvic inflammatory disease. Am J Obstet Gynecol. 2021;224:251–7. 10.1016/j.ajog.2020.10.019.

5. Morrill S, Gilbert NM, Lewis AL. Gardnerella vaginalis as a Cause of Bacterial Vaginosis: Appraisal of the Evidence From in vivo Models. Front Cell Infect Microbiol. 2020;10:168. 10.3389/fcimb.2020.00168.

6. Hickey RJ, Forney LJ. Gardnerella vaginalis does not always cause Bacterial Vaginosis. J Infect Dis. 2014;210:1682–3. 10.1093/infdis/jiu303.

7. Chen X, Lu Y, Chen T, Li R. The Female Vaginal Microbiome in Health and Bacterial Vaginosis. Front Cell Infect Microbiol. 2021;11:631972. 10.3389/fcimb.2021.631972.

8. Hardy L, Jespers V, Abdellati S, De Baetselier I, Mwambarangwe L, Musengamana V, et al. A fruitful alliance: the synergy between Atopobium vaginae and Gardnerella vaginalis in bacterial vaginosis-associated biofilm. Sex Transm Infect. 2016;92:487–91. 10.1136/sextrans-2015-052475.

9. George SD, Amerson-Brown MH, Sousa LGV, Rinehart AH, Tamhane A, Riegler AN, et al. Spatial organization of Gardnerella species, Prevotella bivia, and Fannyhessea vaginae in the bacterial vaginosis biofilm. Infect Immun. 2026;94:e00630–25. 10.1128/iai.00630-25.

10. Tarracchini C, Lugli GA, Mancabelli L, Milani C, Turroni F, Ventura M. Assessing the Genomic Variability of Gardnerella vaginalis through Comparative Genomic Analyses: Evolutionary and Ecological Implications. Appl Environ Microbiol. 2020;87:e02188–20. 10.1128/AEM.02188-20.

11. Harwich MD, Alves JM, Buck GA, Strauss JF, Patterson JL, Oki AT, et al. Drawing the line between commensal and pathogenic Gardnerella vaginalis through genome analysis and virulence studies. BMC Genomics. 2010;11:375. 10.1186/1471-2164-11-375.

12. Jung H-S, Ehlers MM, Lombaard H, Redelinghuys MJ, Kock MM. Etiology of bacterial vaginosis and polymicrobial biofilm formation. Crit Rev Microbiol. 2017;43:651–67. 10.1080/1040841X.2017.1291579.

13. Vaneechoutte M, Guschin A, Van Simaey L, Gansemans Y, Van Nieuwerburgh F, Cools P. Emended description of Gardnerella vaginalis and description of Gardnerella leopoldii sp. nov., Gardnerella piotii sp. nov. and Gardnerella swidsinskii sp. nov., with delineation of 13 genomic species within the genus Gardnerella. Int J Syst Evol Microbiol. 2019;69:679–87. 10.1099/ijsem.0.003200.

14. Verstraelen H, Swidsinski A. The biofilm in bacterial vaginosis: implications for epidemiology, diagnosis and treatment. Curr Opin Infect Dis. 2013;26:86–9. 10.1097/QCO.0b013e32835c20cd.

15. Kalia N, Singh J, Kaur M. Microbiota in vaginal health and pathogenesis of recurrent vulvovaginal infections: a critical review. Ann Clin Microbiol Antimicrob. 2020;19:5. 10.1186/s12941-020-0347-4.

16. Srinivasan S, Morgan MT, Fiedler TL, Djukovic D, Hoffman NG, Raftery D, et al. Metabolic Signatures of Bacterial Vaginosis. mBio. 2015;6:e00204–15. 10.1128/mBio.00204-15.

17. Tajadura-Ortega V, Chai W, Roberts LA, Zhang Y, Di Maio A, Decout AC, et al. Identification and characterisation of vaginal bacteria-glycan interactions implicated in reproductive tract health and pregnancy outcomes. Nat Commun. 2025;16:5207. 10.1038/s41467-025-60404-1.

18. Whelan F, Lafita A, Griffiths SC, Cooper REM, Whittingham JL, Turkenburg JP, et al. Defining the remarkable structural malleability of a bacterial surface protein Rib domain implicated in infection. Proc Natl Acad Sci. 2019;116:26540–8. 10.1073/pnas.1911776116.

19. Swidsinski A, Doerffel Y, Loening-Baucke V, Swidsinski S, Verstraelen H, Vaneechoutte M, et al. Gardnerella Biofilm Involves Females and Males and Is Transmitted Sexually. Gynecol Obstet Invest. 2010;70:256–63. 10.1159/000314015.

20. Petrova MI, Reid G, Vaneechoutte M, Lebeer S. Lactobacillus inersl]: Friend or Foe? Trends Microbiol. 2017;25:182–91. 10.1016/j.tim.2016.11.007.

21. Srinivasan S, Hoffman NG, Morgan MT, Matsen FA, Fiedler TL, Hall RW, et al. Bacterial Communities in Women with Bacterial Vaginosis: High Resolution Phylogenetic Analyses Reveal Relationships of Microbiota to Clinical Criteria. PLoS ONE. 2012;7:e37818. 10.1371/journal.pone.0037818.

22. Gajer P, Brotman RM, Bai G, Sakamoto J, Schütte UME, Zhong X, et al. Temporal Dynamics of the Human Vaginal Microbiota. Sci Transl Med. 2012;4. 10.1126/scitranslmed.3003605.

23. How SS, Nathan S, Lam SD, Chieng S. ATP-binding cassette (ABC) transporters: structures and roles in bacterial pathogenesis. J Zhejiang Univ-Sci B. 2025;26:58–75. 10.1631/jzus.B2300641.

24. Berntsson RP-A, Smits SHJ, Schmitt L, Slotboom D-J, Poolman B. A structural classification of substrate-binding proteins. FEBS Lett. 2010;584:2606–17. 10.1016/j.febslet.2010.04.043.

25. Sánchez-Alba L, Varejão N, Durand A, García-Pardo J, Carreras-Caballé M, Amador V, et al. Structural basis for transcriptional regulation by the cell division regulator MraZ in Mycoplasma genitalium. Nat Commun. 2026;17:2132. 10.1038/s41467-026-68809-2.

26. Yen IY, Whitfield GB, Rubinstein JL, Burrows LL, Brun YV, Howell PL. Conformational changes in the motor ATPase CpaF facilitate a rotary mechanism of Tad pilus assembly. Nat Commun. 2025;16:3839. 10.1038/s41467-025-59009-5.

27. Pelicic V. Mechanism of assembly of type 4 filaments: everything you always wanted to know (but were afraid to ask): This article is part of the Bacterial Cell Envelopes collection. Microbiology. 2023;169. 10.1099/mic.0.001311.

28. Whitfield GB, Brun YV. The type IVc pilus: just a Tad different. Curr Opin Microbiol. 2024;79:102468. 10.1016/j.mib.2024.102468.

29. Cheng L, Correia MSP, Higdon SM, Romero Garcia F, Tsiara I, Joffré E, et al. The protective role of commensal gut microbes and their metabolites against bacterial pathogens. Gut Microbes. 2024;16:2356275. 10.1080/19490976.2024.2356275.

30. Yeoman CJ, Yildirim S, Thomas SM, Durkin AS, Torralba M, Sutton G, et al. Comparative Genomics of Gardnerella vaginalis Strains Reveals Substantial Differences in Metabolic and Virulence Potential. PLoS ONE. 2010;5:e12411. 10.1371/journal.pone.0012411.

31. Bohr LL, Mortimer TD, Pepperell CS. Lateral Gene Transfer Shapes Diversity of Gardnerella spp. Front Cell Infect Microbiol. 2020;10:293. 10.3389/fcimb.2020.00293.

32. Innamorati KA, Earl JP, Barrera SC, Ehrlich RL, Aiyeku J, Gordon A, et al. Metronidazole response profiles of Gardnerella species are congruent with phylogenetic and comparative genomic analyses. Genome Med. 2025;17:28. 10.1186/s13073-025-01446-4.

33. Munch MM, Strenk SM, Srinivasan S, Fiedler TL, Proll S, Fredricks DN. Gardnerella Species and Their Association With Bacterial Vaginosis. J Infect Dis. 2024;230:e171–81. 10.1093/infdis/jiae026.

34. Hill JE, Albert AYK, the VOGUE Research Group. Resolution and Cooccurrence Patterns of Gardnerella leopoldii, G. swidsinskii, G. piotii, and G. vaginalis within the Vaginal Microbiome. Infect Immun. 2019;87:e00532–19. 10.1128/IAI.00532-19.

35. Castro J, Machado D, Cerca N. Unveiling the role of Gardnerella vaginalis in polymicrobial Bacterial Vaginosis biofilms: the impact of other vaginal pathogens living as neighbors. ISME J. 2019;13:1306–17. 10.1038/s41396-018-0337-0.

36. Castro J, França A, Bradwell KR, Serrano MG, Jefferson KK, Cerca N. Comparative transcriptomic analysis of Gardnerella vaginalis biofilms vs. planktonic cultures using RNA-seq. Npj Biofilms Microbiomes. 2017;3:3. 10.1038/s41522-017-0012-7.

37. Sousa LGV, Novak J, França A, Muzny CA, Cerca N. Gardnerella vaginalis, Fannyhessea vaginae, and Prevotella bivia Strongly Influence Each Other’s Transcriptome in Triple-Species Biofilms. Microb Ecol. 2024;87:117. 10.1007/s00248-024-02433-9.

38. Johnson AP, Davies HA. Demonstration by electron microscopy of pili on Gardnerella vaginalis. Sex Transm Infect. 1984;60:396–7. 10.1136/sti.60.6.396.

39. Boustouller YL, Johnson AP, Taylor-Robinson D. Pili on Gardnerella vaginalis studied by electronmicroscopy. J Med Microbiol. 1987;23:327–9. 10.1099/00222615-23-4-327.

40. Mom J, Chouikha I, Valette O, Pieulle L, Pelicic V. Systematic functional analysis of the Com pilus in Streptococcus sanguinisl]: a minimalistic type 4 filament dedicated to DNA uptake in monoderm bacteria. mBio. 2024;15:e02667–23. 10.1128/mbio.02667-23.

41. Raynaud C, Sheppard D, Berry J-L, Gurung I, Pelicic V. PilB from Streptococcus sanguinis is a bimodular type IV pilin with a direct role in adhesion. Proc Natl Acad Sci. 2021;118:e2102092118. 10.1073/pnas.2102092118.

42. Pelicic V. Type IV pili: e pluribus unuml]? Mol Microbiol. 2008;68:827–37. 10.1111/j.1365-2958.2008.06197.x.

43. Mishra G, Gupta K, Mohanty S, Mitra S, Jena S. Evaluation of various diagnostic modalities for detection of bacterial vaginosis and aerobic vaginitis: An underdiagnosed entity among women of the reproductive age group. Indian J Pathol Microbiol. 2025;68:562–7. 10.4103/ijpm.ijpm_459_24.

44. Challa A, Sood S, Kachhawa G, Upadhyay AD, Dwivedi SN, Gupta S. Diagnostic concordance between Amsel’s criteria and the Nugent scoring method in the assessment of bacterial vaginosis. Sex Health. 2022;18:512–4. 10.1071/SH21149.

45. Sha BE, Chen HY, Wang QJ, Zariffard MR, Cohen MH, Spear GT. Utility of Amsel Criteria, Nugent Score, and Quantitative PCR for Gardnerella vaginalis, Mycoplasma hominis, and Lactobacillus spp. for Diagnosis of Bacterial Vaginosis in Human Immunodeficiency Virus-Infected Women. J Clin Microbiol. 2005;43:4607–12. 10.1128/JCM.43.9.4607-4612.2005.

46. Marín E, Haesaert A, Padilla L, Adán J, Hernáez ML, Monteoliva L, et al. Unraveling Gardnerella vaginalis Surface Proteins Using Cell Shaving Proteomics. Front Microbiol. 2018;9:975. 10.3389/fmicb.2018.00975.

47. Zhang K, Lu M, Qiu Y, Zhu X, Wang H, Huang Y, et al. Transcriptomic and Proteomic Analysis of Gardnerella vaginalis Responding to Acidic pH and Hydrogen Peroxide Stress. Microorganisms. 2023;11:695. 10.3390/microorganisms11030695.

48. Borgdorff H, Armstrong SD, Tytgat HLP, Xia D, Ndayisaba GF, Wastling JM, et al. Unique Insights in the Cervicovaginal Lactobacillus iners and L. crispatus Proteomes and Their Associations with Microbiota Dysbiosis. PLOS ONE. 2016;11:e0150767. 10.1371/journal.pone.0150767.

49. Al-Madboly LA, Aboulmagd A, El-Salam MA, Kushkevych I, El-Morsi RM. Microbial enzymes as powerful natural anti-biofilm candidates. Microb Cell Factories. 2024;23:343. 10.1186/s12934-024-02610-y.

50. Dillard LR, Glass EM, Lewis AL, Thomas-White K, Papin JA. Metabolic Network Models of the Gardnerella Pangenome Identify Key Interactions with the Vaginal Environment. mSystems. 2023;8:e00689–22. 10.1128/msystems.00689-22.

51. Ceccarani C, Foschi C, Parolin C, D’Antuono A, Gaspari V, Consolandi C, et al. Diversity of vaginal microbiome and metabolome during genital infections. Sci Rep. 2019;9:14095. 10.1038/s41598-019-50410-x.

52. Sakamoto T, Otokawa T, Kono R, Shigeri Y, Watanabe K. A C69-family cysteine dipeptidase from Lactobacillus farciminis JCM1097 possesses strong Gly-Pro hydrolytic activity. J Biochem (Tokyo). 2013;154:419–27. 10.1093/jb/mvt069.

53. Varmanen P, Savijoki K, Åvall S, Palva A, Tynkkynen S. X-Prolyl Dipeptidyl Aminopeptidase Gene (pepX) Is Part of the glnRA Operon in Lactobacillus rhamnosus. J Bacteriol. 2000;182:146–54. 10.1128/JB.182.1.146-154.2000.

54. Zhang J, Li L, Dan W, Li J, Zhang Q, Bai H, et al. Synthesis and Antimicrobial Activities of 3-Methyl-β-Carboline Derivatives. Nat Prod Commun. 2015;10:899–902.

55. Carstens H, Kalka K, Verhaegh R, Schumacher F, Soddemann M, Wilker B, et al. Antimicrobial effects of inhaled sphingosine against Pseudomonas aeruginosa in isolated ventilated and perfused pig lungs. PLOS ONE. 2022;17:e0271620. 10.1371/journal.pone.0271620.

56. Cheng L, Kaźmierczak D, Norenhag J, Hamsten M, Fransson E, Schuppe-Koistinen I, et al. A MicroRNA Gene Panel Predicts the Vaginal Microbiota Composition. mSystems. 2021;6:e00175–21. 10.1128/mSystems.00175-21.

57. Cheng L, Norenhag J, Hu YOO, Brusselaers N, Fransson E, Ährlund-Richter A, et al. Vaginal microbiota and human papillomavirus infection among young Swedish women. Npj Biofilms Microbiomes. 2020;6:39. 10.1038/s41522-020-00146-8.

58. Ährlund-Richter A, Cheng L, Hu YOO, Svensson M, Pennhag AAL, Ursu RG, et al. Changes in Cervical Human Papillomavirus (HPV) Prevalence at a Youth Clinic in Stockholm, Sweden, a Decade After the Introduction of the HPV Vaccine. Front Cell Infect Microbiol. 2019;9:59. 10.3389/fcimb.2019.00059.

59. De Coster W, Rademakers R. NanoPack2: population-scale evaluation of long-read sequencing data. Bioinformatics. 2023;39:btad311. 10.1093/bioinformatics/btad311.

60. Kolmogorov M, Yuan J, Lin Y, Pevzner PA. Assembly of long, error-prone reads using repeat graphs. Nat Biotechnol. 2019;37:540–6. 10.1038/s41587-019-0072-8.

61. Hackl T, Trigodet F, Eren AM, Biller SJ, Eppley JM, Luo E, et al. proovframe: frameshift-correction for long-read (meta)genomics. 2021. 10.1101/2021.08.23.457338.

62. Buchfink B, Reuter K, Drost H-G. Sensitive protein alignments at tree-of-life scale using DIAMOND. Nat Methods. 2021;18:366–8. 10.1038/s41592-021-01101-x.

63. Sereika M, Kirkegaard RH, Karst SM, Michaelsen TY, Sørensen EA, Wollenberg RD, et al. Oxford Nanopore R10.4 long-read sequencing enables the generation of near-finished bacterial genomes from pure cultures and metagenomes without short-read or reference polishing. Nat Methods. 2022;19:823–6. 10.1038/s41592-022-01539-7.

64. Parks DH, Imelfort M, Skennerton CT, Hugenholtz P, Tyson GW. CheckM: assessing the quality of microbial genomes recovered from isolates, single cells, and metagenomes. Genome Res. 2015;25:1043–55. 10.1101/gr.186072.114.

65. Tatusova T, DiCuccio M, Badretdin A, Chetvernin V, Nawrocki EP, Zaslavsky L, et al. NCBI prokaryotic genome annotation pipeline. Nucleic Acids Res. 2016;44:6614–24. 10.1093/nar/gkw569.

66. Gautreau G, Bazin A, Gachet M, Planel R, Burlot L, Dubois M, et al. PPanGGOLiN: Depicting microbial diversity via a partitioned pangenome graph. PLOS Comput Biol. 2020;16:e1007732. 10.1371/journal.pcbi.1007732.

67. Bastian M, Heymann S, Jacomy M. Gephi: An Open Source Software for Exploring and Manipulating Networks. Proc Int AAAI Conf Web Soc Media. 2009;3:361–2. 10.1609/icwsm.v3i1.13937.

68. Bazin A, Medigue C, Vallenet D, Calteau A. panModule: detecting conserved modules in the variable regions of a pangenome graph. 2021. 10.1101/2021.12.06.471380.

69. Szklarczyk D, Nastou K, Koutrouli M, Kirsch R, Mehryary F, Hachilif R, et al. The STRING database in 2025: protein networks with directionality of regulation. Nucleic Acids Res. 2025;53:D730–7. 10.1093/nar/gkae1113.

70. Abramson J, Adler J, Dunger J, Evans R, Green T, Pritzel A, et al. Accurate structure prediction of biomolecular interactions with AlphaFold 3. Nature. 2024;630:493–500. 10.1038/s41586-024-07487-w.

71. Hallgren J, Tsirigos KD, Pedersen MD, Almagro Armenteros JJ, Marcatili P, Nielsen H, et al. DeepTMHMM predicts alpha and beta transmembrane proteins using deep neural networks. 2022. 10.1101/2022.04.08.487609.

72. Blum M, Andreeva A, Florentino LC, Chuguransky SR, Grego T, Hobbs E, et al. InterPro: the protein sequence classification resource in 2025. Nucleic Acids Res. 2025;53:D444–56. 10.1093/nar/gkae1082.

73. Stoddard SF, Smith BJ, Hein R, Roller BRK, Schmidt TM. rrnDB: improved tools for interpreting rRNA gene abundance in bacteria and archaea and a new foundation for future development. Nucleic Acids Res. 2015;43:D593–8. 10.1093/nar/gku1201.

74. Gao Y, Wu M. Accounting for 16S rRNA copy number prediction uncertainty and its implications in bacterial diversity analyses. ISME Commun. 2023;3:59. 10.1038/s43705-023-00266-0.

75. Tautenhahn R, Patti GJ, Rinehart D, Siuzdak G. XCMS Online: A Web-Based Platform to Process Untargeted Metabolomic Data. Anal Chem. 2012;84:5035–9. 10.1021/ac300698c.

76. Forsberg EM, Huan T, Rinehart D, Benton HP, Warth B, Hilmers B, et al. Data processing, multi-omic pathway mapping, and metabolite activity analysis using XCMS Online. Nat Protoc. 2018;13:633–51. 10.1038/nprot.2017.151.

77. Fan S, Kind T, Cajka T, Hazen SL, Tang WHW, Kaddurah-Daouk R, et al. Systematic Error Removal Using Random Forest for Normalizing Large-Scale Untargeted Lipidomics Data. Anal Chem. 2019;91:3590–6. 10.1021/acs.analchem.8b05592.

78. Wishart DS, Guo A, Oler E, Wang F, Anjum A, Peters H, et al. HMDB 5.0: the Human Metabolome Database for 2022. Nucleic Acids Res. 2022;50:D622–31. 10.1093/nar/gkab1062.

79. Dührkop K, Shen H, Meusel M, Rousu J, Böcker S. Searching molecular structure databases with tandem mass spectra using CSI:FingerID. Proc Natl Acad Sci. 2015;112:12580–5. 10.1073/pnas.1509788112.

80. Hoffmann MA, Nothias L-F, Ludwig M, Fleischauer M, Gentry EC, Witting M, et al. Assigning confidence to structural annotations from mass spectra with COSMIC. 2021. 10.1101/2021.03.18.435634.

81. Rainer J, Vicini A, Salzer L, Stanstrup J, Badia JM, Neumann S, et al. A Modular and Expandable Ecosystem for Metabolomics Data Annotation in R. Metabolites. 2022;12:173. 10.3390/metabo12020173.

82. Horai H, Arita M, Kanaya S, Nihei Y, Ikeda T, Suwa K, et al. MassBank: a public repository for sharing mass spectral data for life sciences. J Mass Spectrom. 2010;45:703–14. 10.1002/jms.1777.

83. Schymanski EL, Jeon J, Gulde R, Fenner K, Ruff M, Singer HP, et al. Identifying Small Molecules via High Resolution Mass Spectrometry: Communicating Confidence. Environ Sci Technol. 2014;48:2097–8. 10.1021/es5002105.

84. Smith TP, Clegg T, Ransome E, Martin-Lilley T, Rosindell J, Woodward G, et al. High-throughput characterization of bacterial responses to complex mixtures of chemical pollutants. Nat Microbiol. 2024;9:938–48. 10.1038/s41564-024-01626-9.

85. Perez-Riverol Y, Bandla C, Kundu DJ, Kamatchinathan S, Bai J, Hewapathirana S, et al. The PRIDE database at 20 years: 2025 update. Nucleic Acids Res. 2025;53:D543–53. 10.1093/nar/gkae1011.

