## Supplementary Data 01 for "Multi-omics uncover interactions in the vaginal microbiome and a type II secretion/Tad pilus system in *Gardnerella* species"

**Supplementary Data S**

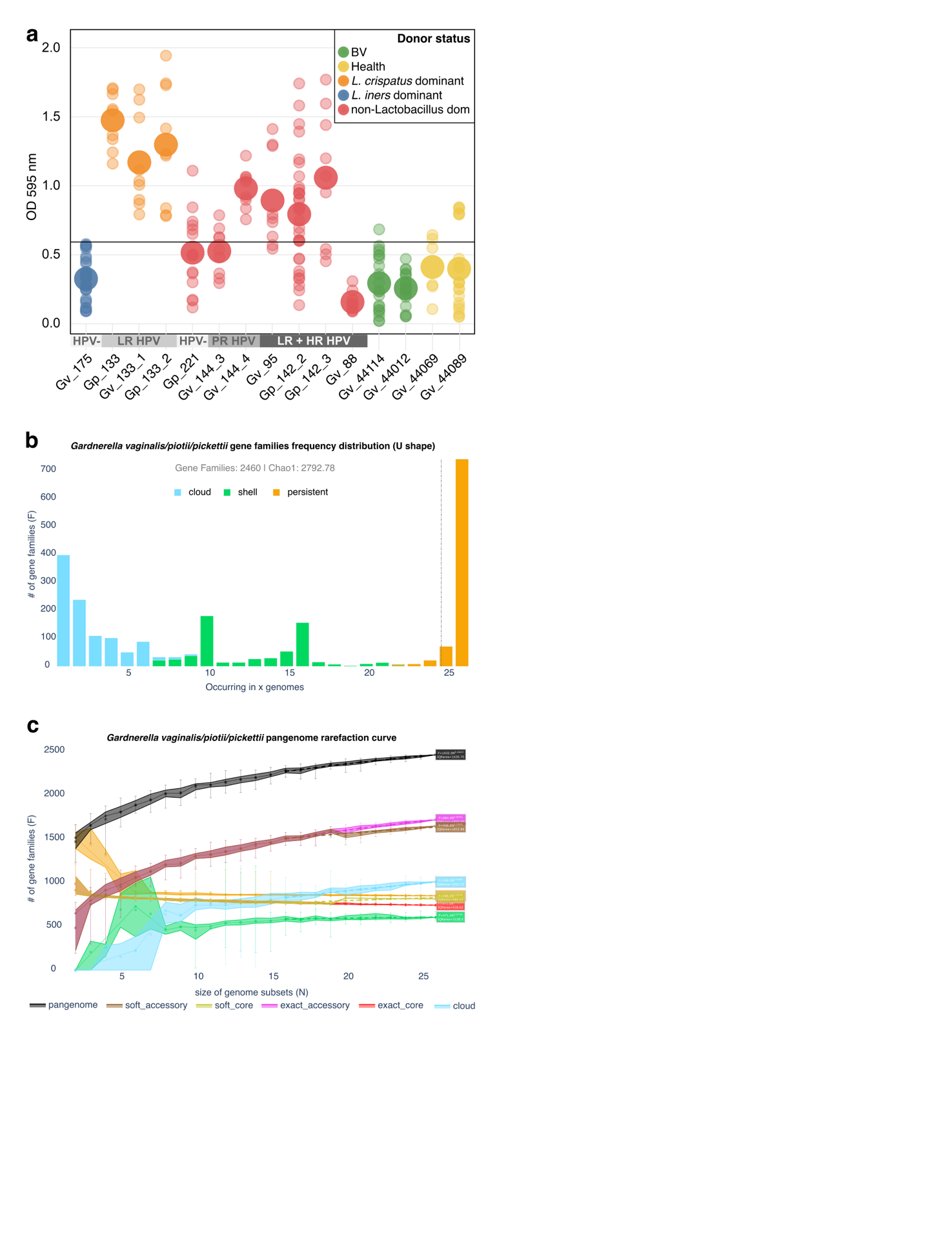

**Fig. S1.** **Scatterplot, U-shaped plots and rarefaction curves/parameters of *Gardnerella* pangenome.** a. Scatterplot showing biofilm formation capacity of 15 *Gardnerella* strains measured by crystal violet assays. Black horizontal line indicates the overall mean (OD_595_ = 0.61). Samples were categorized as having high, medium, or low biofilm formation based on whether their mean value was above, overlapping (overall mean ± 0.15), or below the overall mean, respectively. (n = 3 - 11). b) U-shaped plot of *Gardnerella vaginalis/piotii/pickettii* strains. c) Rarefaction curves of *Gardnerella vaginalis/piotii/pickettii* pangenome. Abbreviations: BV = Bacterial vaginosis, HPV- = human papillomavirus negative, LR HPV = low-risk type human papillomavirus, PR HPV = probably-risk type human papillomavirus, HR HPV = High-risk type human papillomavirus, *Gv* = *Gardnerella vaginalis, Gp* = *Gardnerella piotii/pickettii*.

| *Gardnerella* species rarefaction parameters | | | |  |  |
| --- | --- | --- | --- | --- | --- |
| partition | **kappa** | **gamma** | **kappa_std_error** | **gamma_std_error** | **IQR_area** |
| persistent | 932.11321 | -0.0284618 | 7.45326913 | 0.00263421 | 1376.75 |
| shell | 470.891752 | 0.07734434 | 23.9835061 | 0.01675188 | 3128.5 |
| cloud | 267.609326 | 0.40685436 | 14.647106 | 0.01791329 | 3293.75 |
| exact_core | 971.155831 | -0.082686 | 13.7799902 | 0.00467838 | 518.625 |
| exact_accessory | 683.846868 | 0.28315991 | 15.0336928 | 0.00720822 | 1855.375 |
| soft_core | 495.533132 | 0.1574374 | 12.302619 | 0.0081558 | 486.625 |
| soft_accessory | 936.492628 | 0.17074655 | 22.9795957 | 0.00805921 | 1833.875 |
| pangenome | 1431.88415 | 0.16623446 | 19.6703166 | 0.00451219 | 1630.75 |

**Table S2.- Rarefaction parameters of *Gardnerella* pangenome.**

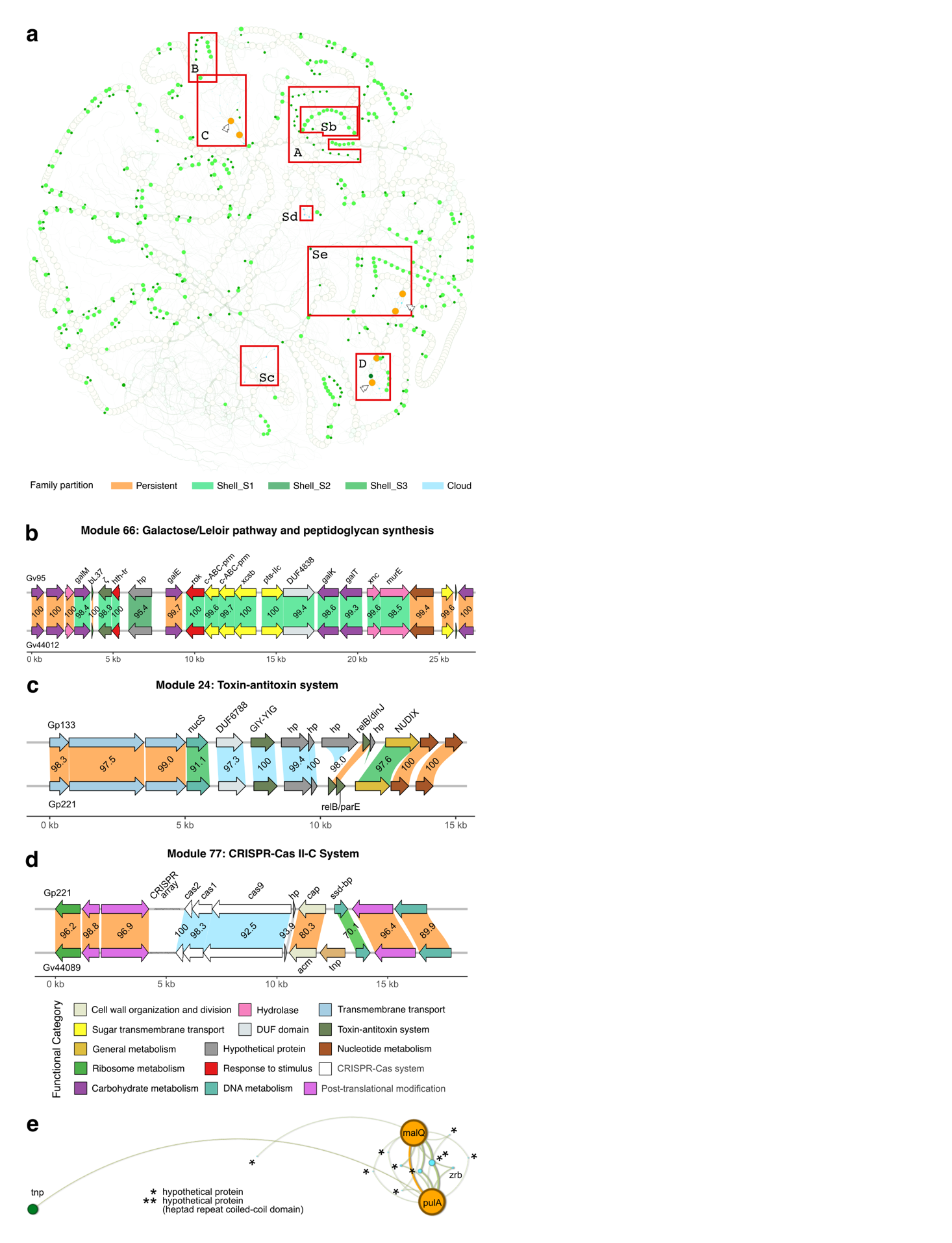

**Fig. S2.** **Syntenic network, gene map and hotspot locus of *Gardnerella* pangenome.** a) Syntenic network displaying genomic islands and divergent regions of conserved modules present in the accessory genome. Circles represent gene families, while edges represent genomic neighbouring. Circle size represents the number of genomes in which the gene family is represented. Examples of hypervariable gene hotspot loci are indicated with white arrows. Enclosed in red are genomic modules or hotspot loci shown in the panels labelled accordingly. b) Module 66: Galactose/Leloir pathway and peptidoglycan synthesis. c) Module 24: Toxin-antitoxin system. d) Module 77: CRISPR-Cas II-C system. For additional genomic context, each module is depicted with three non-module gene families flanking its upstream and downstream regions. The vertical linking regions between synteny maps are coloured by pangenomic partition. Percentage amino acid identity between homologs is shown above each linking area. Network diagram depicts hypervariable gene families converging on core gene families, with the latter serving as gravitational centres; e) hypothetical protein (*) and hypothetical protein (heptad repeat coiled-coil domain) (**) converge on both type 1 pullulanase (pulA) and 4-alpha-glucanotransferase (malQ). Full gene names for abbreviations used in the figure are provided in Supplementary Tables S3.

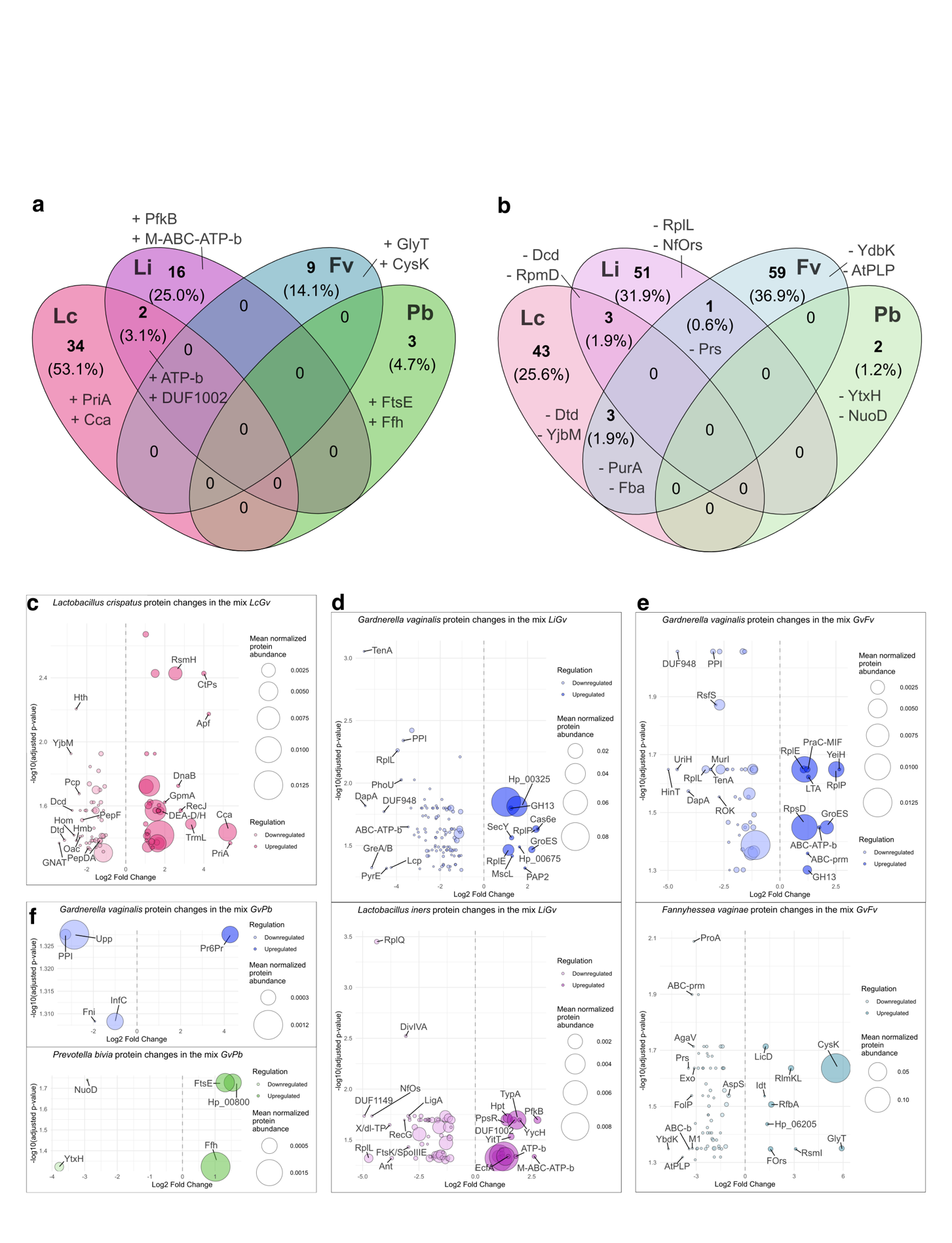

**Fig. S3**. **Species-specific proteomic interactions shape the dynamics of vaginal microbial co-cultures.** a) Venn diagram showing the number and relative percentage of upregulated proteins in *Lactobacillus crispatus* (*Lc*), *Lactobacillus iners* (*Li*), *Fannyhessea vaginae* (*Fv*), and *Prevotella bivia* (*Pb*) when co-cultured with *Gv*. The top two proteins are labelled per condition/overlap, when available. b) Venn diagram showing the number and relative percentage of downregulated proteins in Li, *Fv* and Pb when co-cultured with *Gv.* c) Bubble plot showing *Lc* in the co-culture *LcGv.* Downregulated proteins are on the left side in light colours and upregulated proteins on the right side. Circle size indicates the mean normalized protein abundance. The top 20 most fold-changed proteins are labelled, (FC > 1) adjusted *p-value* (< 0.05) (Welch’s t-test followed by a Benjamin-Hochberg multiple test correction). d) Bubble plot showing *Gv* (top panel) and *Li* (bottom panel) in the co-culture *LiGv.* e) *Gv* (top) and *Fv* (bottom) in the co-culture *GvFv*; and f) *Gv* (top) and *Pb* (bottom) in the co-culture *GvPb*. All conditions n=3 , except for GvPb, which are n=2. Abbreviations: *Lc* = *Lactobacillus crispatus,* *Li* = *Lactobacillus iners*, *Gv* = *Gardnerella vaginalis*, *Fv* = *Fannyhessea vaginae*, and *Pb* = *Prevotella bivia*. Full gene names for abbreviations used in the figure are provided in Supplementary Tables S3.

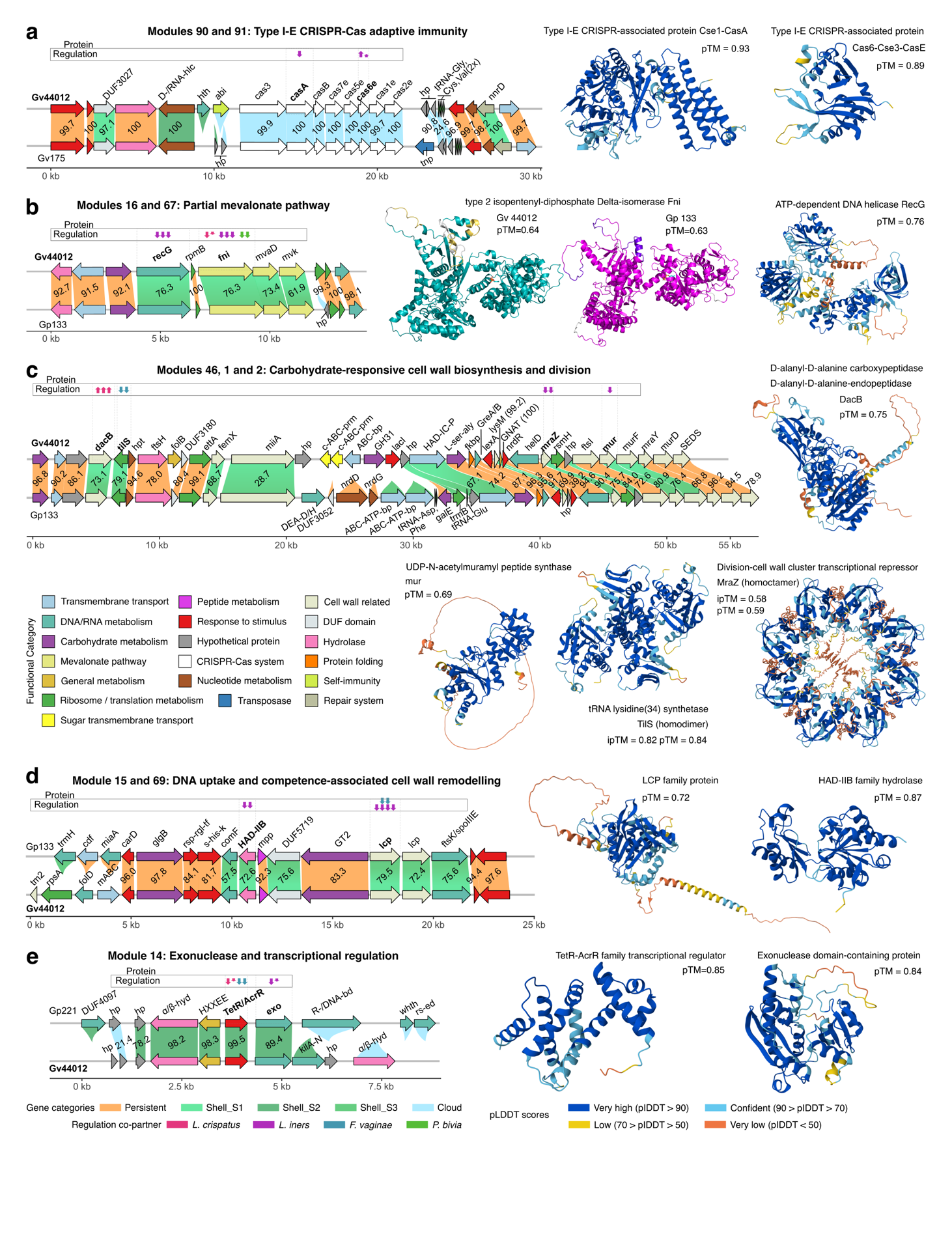

**Fig. S4. Gena maps of regulated Gv proteins and AI predicted structures**. a) Modules 90 and 91: Type I-E CRISPR-Cas adaptive immunity; Protein structures: CasA, Gv44012, pTM = 0.93; Cas6e, Gv44012, pTM = 0.89. b) Modules 16 and 67: Partial mevalonate pathway for isoprenoid biosynthesis; Fni, Gv44012, pTM=0.64; Gp133, pTM=0.63. AI predicted protein structures comparing homologous from *Gv* (cyan coloured structure) and *Gp* (magenta coloured structure). Amino acid sequence divergence is highlighted in yellow and purple in Gv and Gp structures respectively, while in white is highlighted inserted regions in both cases; RecG, Gv44012, pTM = 0.76. c) Modules 46, 1 and 2: Carbohydrate-responsive cell wall biosynthesis and division; DacB, pTM = 0.75; Mur, pTM = 0.69; TilS (homodimer), ipTM = 0.82, pTM = 0.84; MraZ (homoctamer), ipTM = 0.58, pTM = 0.59; d) Module 15 and 69: DNA uptake and competence-associated cell wall remodelling; Lcp, pTM = 0.72; HAD-IIB, pTM = 0.87; e) Module 14: Exonuclease and transcriptional regulation; TetR/AcrR, pTM = 0.85; Exo, pTM = 0.84. For additional genomic context, each gene module is depicted with three non-module gene families flanking its upstream and downstream regions. Protein regulation: Regulated gene names are presented in bold with arrows indicating direction (one arrow = 1.0-1.4 |FC|; one and a half arrows = 1.45-1.85 |FC|, etc). Arrow colors indicate the bacterial species triggering the regulation. Colours in single protein models (non comparisons) appear as defined in Alpha Fold Server, where “pLDDT: a per-atom confidence estimate on a 0-100 scale where a higher value indicates higher confidence”, pTM = predicted template modelling score. Full gene names for abbreviations used in the figure are provided in Supplementary Tables S3.

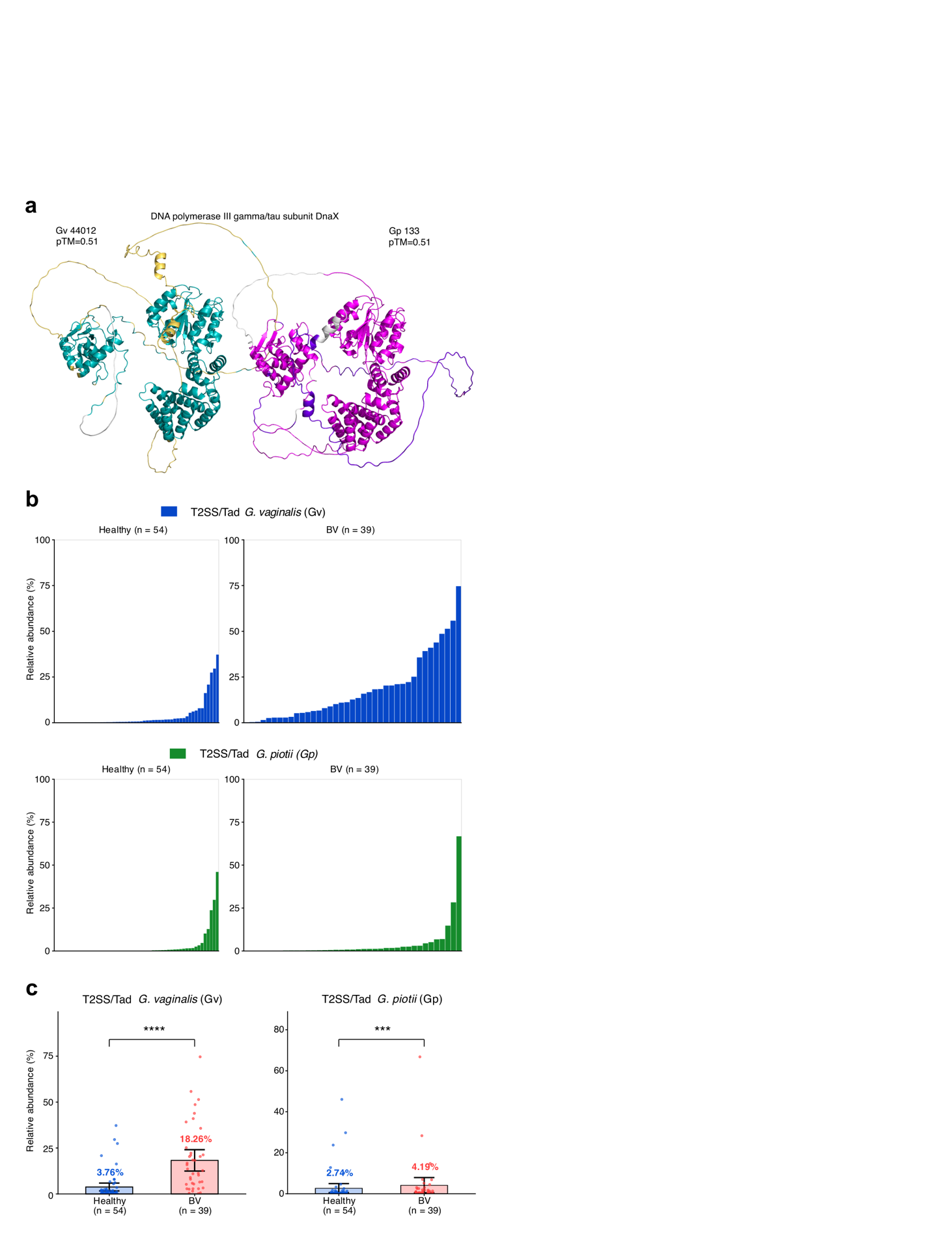

**Fig. S5. DnaX protein predicted structures. Type II secretion/ Tad pilus system (T2SS/Tad) putative operon per-sample relative abundance and per-health status mean.**

a) DNA polymerase III gamma/tau subunit, Gv44012: pTM=0.51; Gp133: pTM=0.51. AI predicted protein structures comparing homologous from *Gv* (cyan coloured structure) and *Gp* (magenta coloured structure). Amino acid sequence divergence is highlighted in yellow and purple in Gv and Gp structures respectively, while in white is highlighted inserted regions in both cases. b) T2SS/Tad putative operon relative abundance per-sample. Healthy samples = 54, BV samples = 39. *Gardnerella vaginalis* = dark blue, *G. piotii* = green; samples are ordered by relative abundance. c) T2SS/Tad putative operon relative abundance per-health status. Healthy = dark blue, BV = Red. Points represent individual samples, bars represent group means, and whiskers indicate 95% confidence intervals of the mean. Statistical significance was assessed using a two-sided Wilcoxon rank-sum test. *** *p* < 0.001; **** *p* < 0.0001. For Gv, *p* = 4.45 × 10⁻⁹; for Gp, *p* = 7.37 × 10⁻⁴.

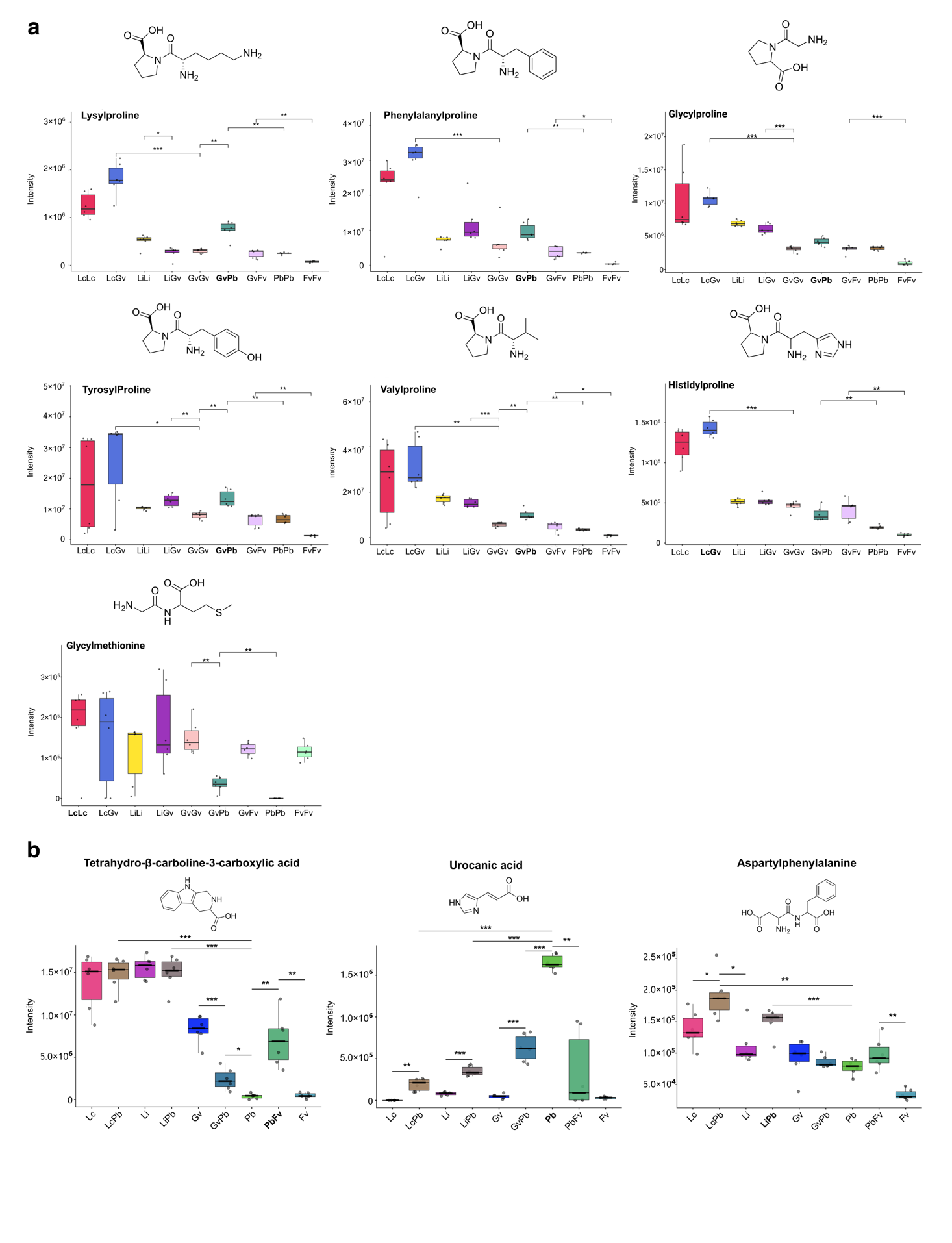

**Fig. S6**. **Additional dipeptides and metabolites at significant high levels in *G. vaginalis* and *P. bivia* cocultures*.*** a, b) Bar plots show intensity distributions for the indicated metabolite across relevant conditions. Individual biological (x3) and technical (x2) replicates are shown as black jittered points. Whiskers extend to the most extreme values within 1.5 x the interquartile range (IQR). The median is marked by a horizontal crossbar. Chemical structures of relevant metabolites are displayed on top of each plot. Statistical comparisons (Welch’s t‑test, FDR‑adjusted) are indicated where applicable (* *p* < 0.05, ** *p* < 0.01, *** *p* < 0.001).

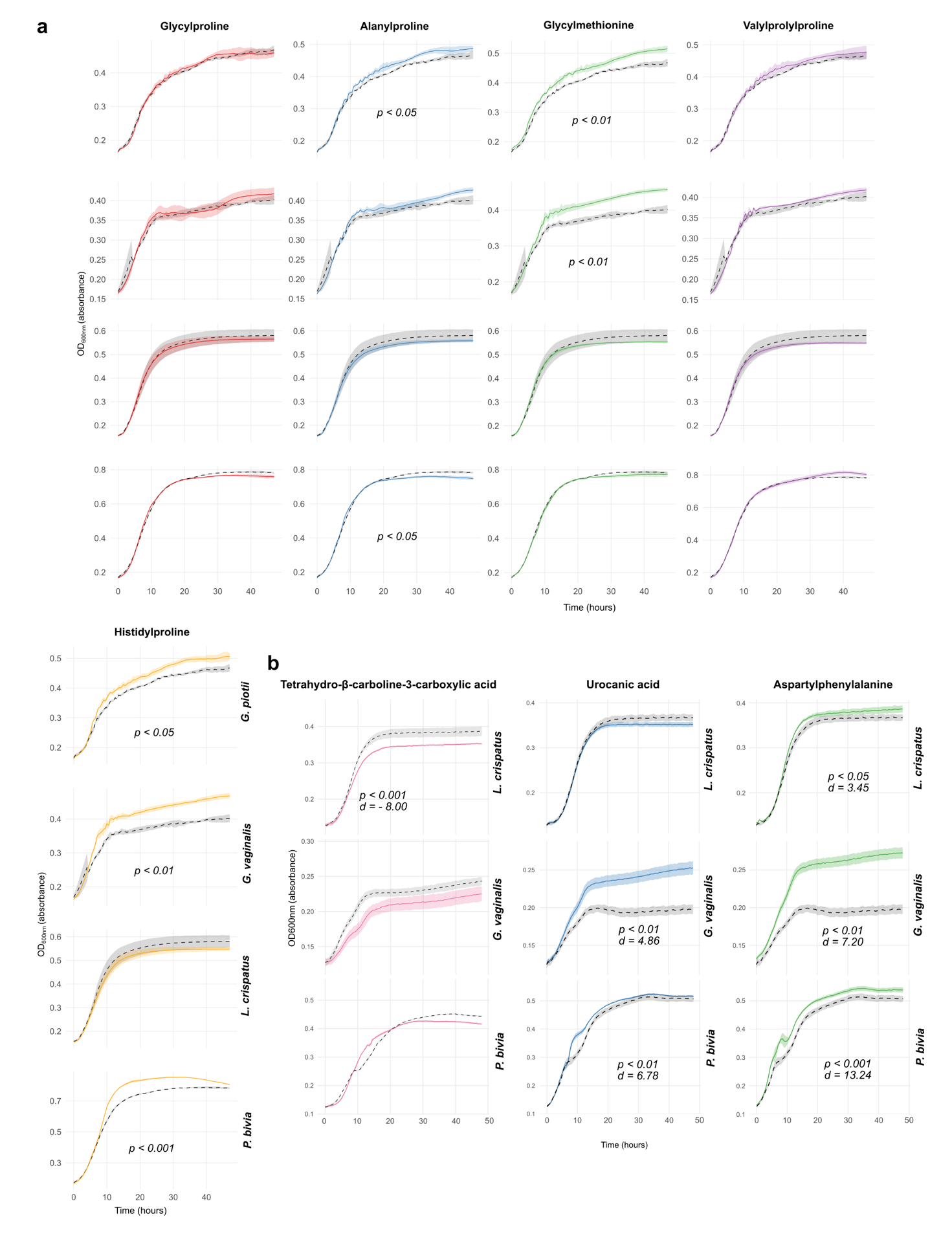

**Fig. S7**. **Bacterial derived proline/methionine/phenylalanine-based dipeptides and tripeptide, and acids effects in vaginal related bacteria*.*** a) Growth curves (Optical density [OD] 600 nm) of relevant vaginal related bacteria cultured with 5.0 mM of different overexpressed metabolites in cocultures containing *Gv*. A single representative experimental curve is shown per condition/strain. Growth curves represent the mean of three biological replicates, each assayed thrice (Gly-Pro and Gly-Met, n = 3) or twice (Ala-Pro, Val-Pro-Pro and His-Pro, n = 2) at 1.0 mM, 5.0 mM or 10 mM with similar inhibitory effects. Alpha-channel-coloured areas correspond to the standard deviation of each experiment. Statistical significance between treatment and control AUC groups is shown using unpaired two-tailed Student's t-tests. gray = medium (control). Abbreviations: AUC = area under the curve. b) Growth curves (Optical density [OD] 600 nm) of relevant vaginal related bacteria cultured with 1.0 mM of different overexpressed metabolites in cocultures containing *Pb*. A single representative experimental curve is shown per condition/strain. Growth curves represent the mean of three biological replicates, each assayed thrice. Alpha-channel-coloured areas correspond to the standard deviation of each experiment. Statistical significance between treatment and control AUC groups is shown using unpaired two-tailed Student's t-tests, with effect sizes reported as Cohen's d.
