## Supplementary information S2, S3 for "Multi-omics uncover interactions in the vaginal microbiome and a type II secretion/Tad pilus system in *Gardnerella* species"

**S2.** ***L. iners* biofilm and pangenome analysis**

***L. iners* show a broad biofilm phenotype and a large continuous syntenic core pangenome**

We assessed biofilm phenotype of 11 *Li* isolates using the same approach applied to *Gardnerella* (Supplementary figure 1a). However, unlike *Gardnerella*, the *Li* genomes clustered according to a distinct phylogenetic logic that did not show a comparable association with biofilm phenotype (Supplementary figure 1b). In addition, the *Li* pangenome displayed a more continuous syntenic core, being less fragmented by accessory elements. Consistent with this, the *Li* pangenome contained a larger core partition (53.56%), followed by cloud (36.96%) and shell (9.48%) (Supplementary figure 1c).

Furthermore, *Li* pangenome revealed genomic islands carrying genes implicated in multiple sugar uptake and metabolism, bacteriophage tail assembly, transcriptional regulation, heavy metal homeostasis, integrase-mediated mobilization, site-specific methylation, and conjugation-associated systems (e.g., ArdA-FtsK), suggesting that these islands contribute to niche adaptation and inter-strain competition (Supplementary figure 1d, 1e).

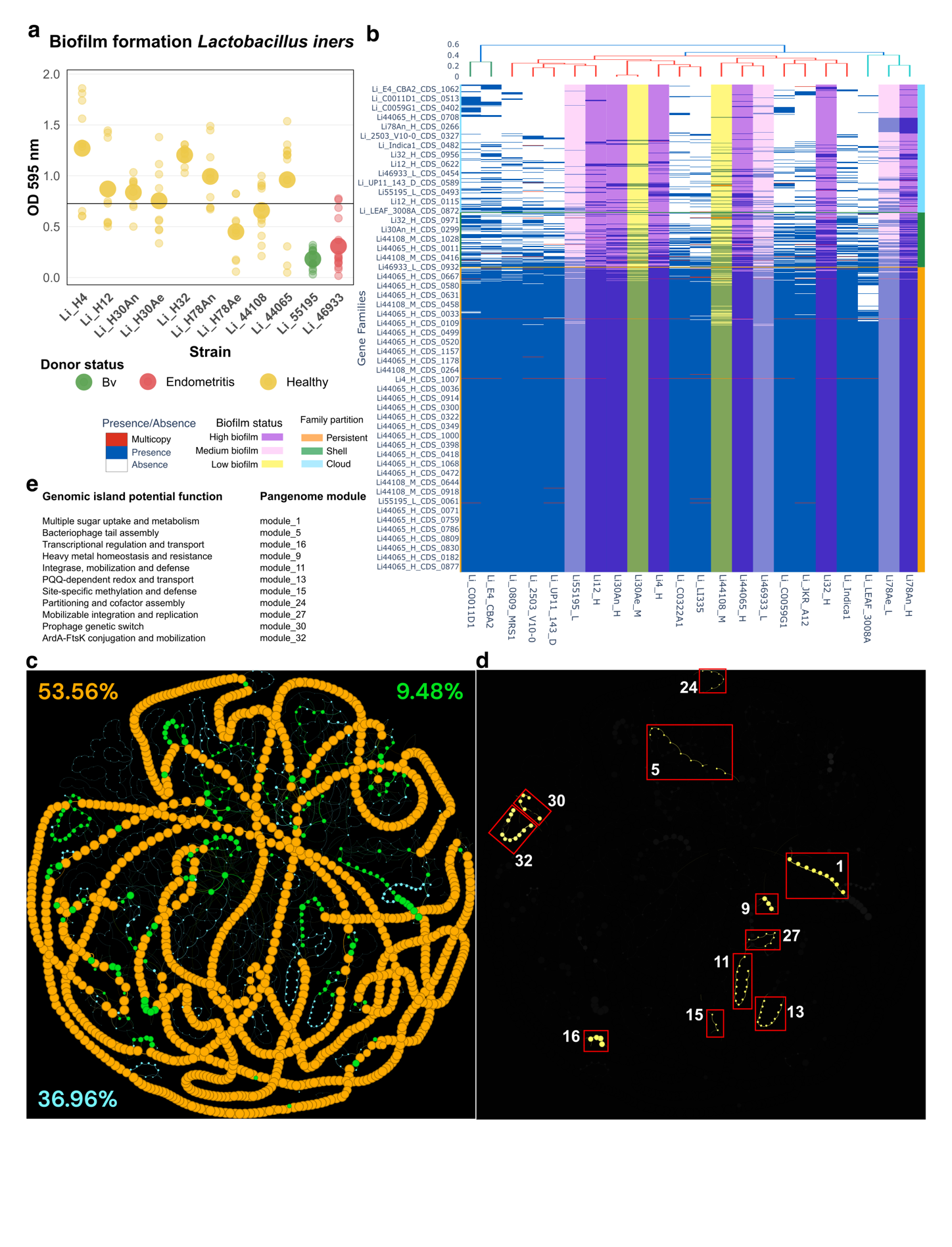

**Supplementary figure 1. Biofilm formation, gene family matrix and syntenic maps of *Lactobacillus iners (Li)*.** a) Scatterplot showing biofilm formation capacity of 11 *Li* strains measured by crystal violet assays. Black horizontal line indicates the overall mean (OD_595_ = 0.73). Samples were categorized as having high, medium, or low biofilm formation based on whether their mean value was above, overlapping, or below the overall mean, respectively (n = 3 – 5). b) Syntenic network mapping gene families onto pangenome partitions in *Li*. Circles represent gene families, while edges represent genomic neighbouring. Size of the circles represent the number of genomes the gene family is represented in. c) Presence-absence matrix of gene families in *Li* genomes. The gene families are ordered by partition and then by their number of presences (increasing order). Genomes are ordered by hierarchical clustering based on their shared gene families via a Jaccard distance. Genomes with biofilm typing are highlighted in coloured transparency; pink = medium biofilm formation, purple = high biofilm formation and yellow = low biofilm formation. d) Syntenic network displaying genomic islands of conserved modules present in accessory genome. Highlighted in yellow are gene families present in high biofilm formation strains. Enclosed in red are genomic islands shown in list e. e) List of potential functional processes of genomic island and module number.

**U-shaped plot and rarefaction curves**

A characteristic U-shaped plot of gene families in *Li* shows a typical distribution in which bacterial gene families tend to be either rare (accessory, lineage-specific) or nearly universal (core, essential), with relatively fewer families at intermediate prevalence (Supplementary Fig. 1a). Rarefaction analysis reveals that *Li* pangenome is open (γ = 0.1181), exhibiting an expanding shell compartment (γ = 0.5526) driving diversity (Supplementary Fig. 1b; Supplementary Table 1).

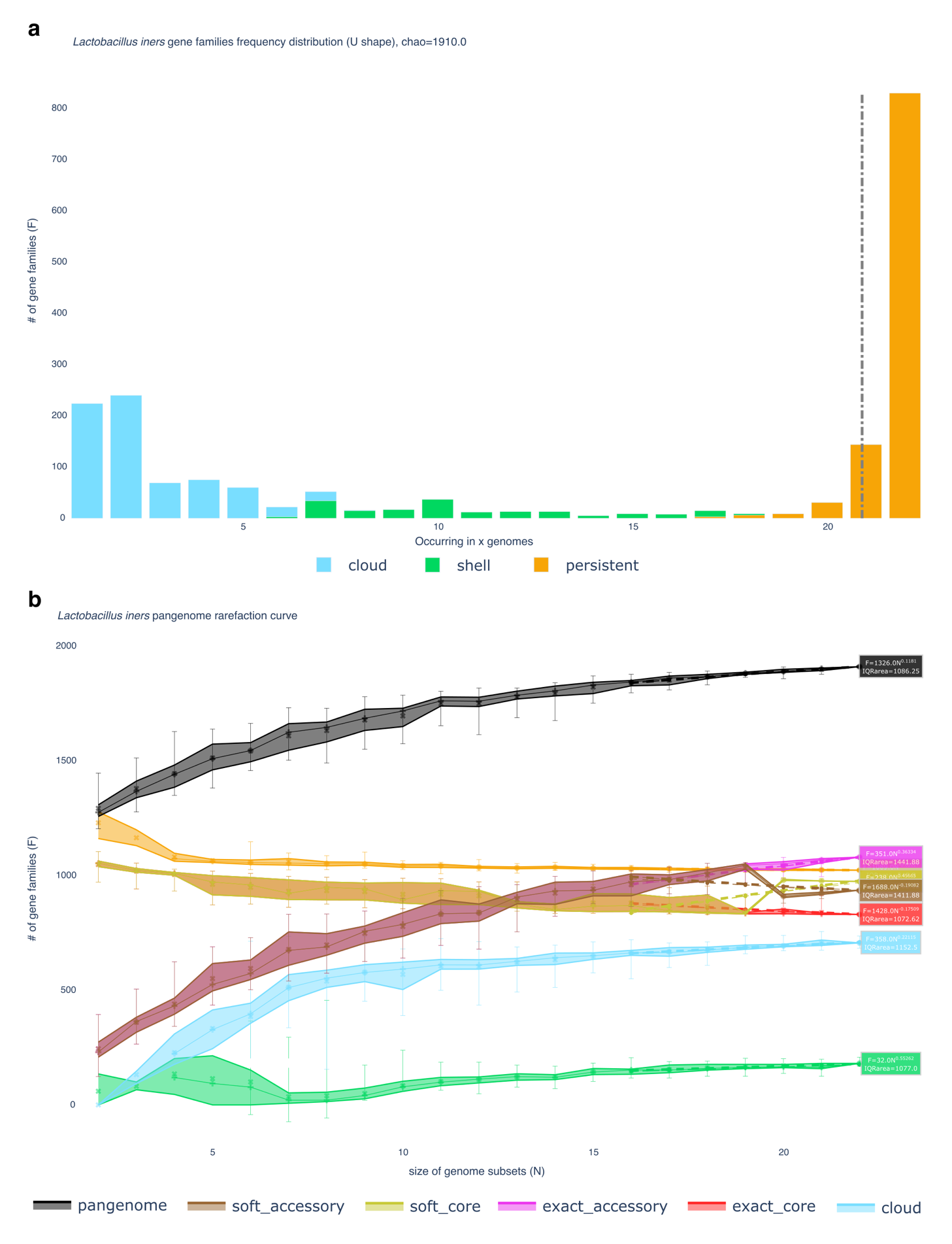

**Supplementary figure 2. U-shaped plots and rarefaction curves of *Gardnerella vaginalis* (*Gv*) and *Lactobacillus iners* (*Li)* pangenomes.** a) U-shaped plots of *Gv* and *Li* penagenomes. b) Rarefaction curves of *Gv* and *Li* penagenomes.

**Supplementary table 1. Rarefaction parameters of *Lactobacillus iners* pangenome. (see file)**

**Supplementary table 2.** **List of modules (conserved regions of genomic plasticity) present in high biofilm in *Lactobacillus iners*. (see file)**

**Supplementary table 3.** **List of all gene families present in *Lactobacillus iners* pangenome. (see file)**

**Supplementary table 4.** **List of all abbreviations for Supplementary Figure 3, 4, 5 and 6 (see file)**

**S3. Additional altered proteins after pairwise co-culture comparisons**

**Species-specific proteomic responses of *Lactobacillus crispatus* (*Lc*) to other vaginal bacteria**

*Lc* mounted a core strategy towards all four vaginal bacteria via upregulation of anchoring and modifying essential cell wall glycopolymers (WaaL, ~2.15), and peptidoglycan recycling and cell wall turnover (MupG, ~1.52) with *Lactobacillus iners* (*Li*), *Gardnerella vaginalis* (*Gv*) and *Fannyhessea vaginae* (*Fv*); auto-/co-aggregation, (Apf, ~4.28) with *Gv* and *Prevotella bivia* (*Pb*); biosynthesis of L-threonine (ThrB, ~1.78) with *Li* and *Fv*; genomic stability maintenance (*RarA*, ~1.95) with *Gv* and *Fv*; cellular defense and environmental adaptation (Dcm, ~1.55) with *Li* and *Gv*. Partner-specific upregulatory adaptations included phospholipid biosynthesis with *Li* (CdsA, 2.57); replication restart, and protein synthesis and maintenace with *Gv* (PriA, 5.35; and *Cca*, 5.22, respectively); membrane transportation with *Fv* (ABC-prm, 2.22), and quorum sensing, and breaking down of proline dense polypeptides to dipeptides with *Pb* (SepM, 4.54; and PepX, 3.56, respectively) (Supplementary Fig. 3a, 4a, 4c, 4e and Figure 3d). These findings demonstrated that *Lc* induced species-specific transcriptional upregulations across cell wall maintenance, metabolism, genomic stability, transport, and quorum sensing pathways in a partner-dependent manner.

*Lc* downregulated probiotic related protein and 16S regulation when co-cultured with *Li*, *Gv* and *Fv* (P8, ~-2.61;RbfA, ~-1.46, respectively), cell wall stabilization (Lcp, ~-1.68) with *Li* and *Fv*; cell wall building and modification (GT4, ~-2.87) with *Gv* and *Pb*; membrane homeostasis (Hly, ~-1.70) with *Li* and *Pb*, stringent response (YjbM, ~-2.90) with *Li* and *Gv*; response regulation (Hth, ~-2.70) with *Gv* and *Fv*; post-transcriptional regulation (S1rb, ~-1.61) with *Fv* and *Pb*. Partner-specific downregulation accounted for cell wall bacterial adhesion with *Li* (CAP, -3.82), translational modulation with *Gv* (Dtd, -3.14), cell membrane remodelling and lipid metabolism with *Fv* (Pld, -3.63), and amino acid uptake and cell-wall modification with *Pb* (Aa-ABC-ATP-b, 2.07; C40, -1.98, respectively) (Supplementary Fig. 3b ,4a, 4c, 4e and Figure 3d). These findings reveal that *Lc* broadly suppresses cell wall dynamics and host interaction pathways, while tailoring downregulation of adherence and metabolic functions to specific bacterial partners.

The presence of *Lc* elicited distinct responses in the *BV*-associated bacteria. Shared upregulation was limited with arginine metabolism regulaton in *Fv* and *Gv* (ArgR, ~3.16); response regulation (Hth, ~-2.83) in *Fv* and *Pb*; and ATP-dependent membrane transport (ABC-ATP-b, ~1.01) in *Gv* and *Pb*. Unique-partner augmented responses included mucin-binding adhesins in *Li* (InlB, 2.13), DNA recombination in *Gv* (Rec, 5.69); iron acquisition coupled to cofactor regulation of metabolism in *Fv* (SufD, 2.87), and nutrient fermentation and anaerobic respiration in *Pb* (CitX, 5.19) (Supplementary Fig. 3c ,4b, 4d, 4f and Figure 3d). Overall, Lc triggered limited shared responses but drove species-specific metabolic adaptations.

Moreover, *Lc* commonly downregulated anaerobic DNA-synthesis/repair in *Gv, Fv* and *Pb* (NrdD, ~-2.11); aromatic amino acids biosynthesis in *Gv* and *Fv* (AroA, ~-1.94); general metabolism in *Fv* and *Pb* (FolC, ~-2.86); and transcription and repair in *Gv* and *Pb* (Mtd, ~-3.50). Species-specific vulnerabilities included lactose utilization (LacB, -2.22) in *Li*; DNA recombination and repair (RuvC, -2.16) in *Gv*; epigenetic regulation (Sdmt, -3.04) in *Fv*; and DNA biosynthesis regulation and complex carbohydrates degradation (NrdJ, -4.29 and GH97, -4.12, respectively) in *Pb*. (Supplementary Fig. 3d ,4b, 4d, 4f and Figure 3d). This showed that Lc modified the biosynthesis functions of other BV-associated bacterial species.

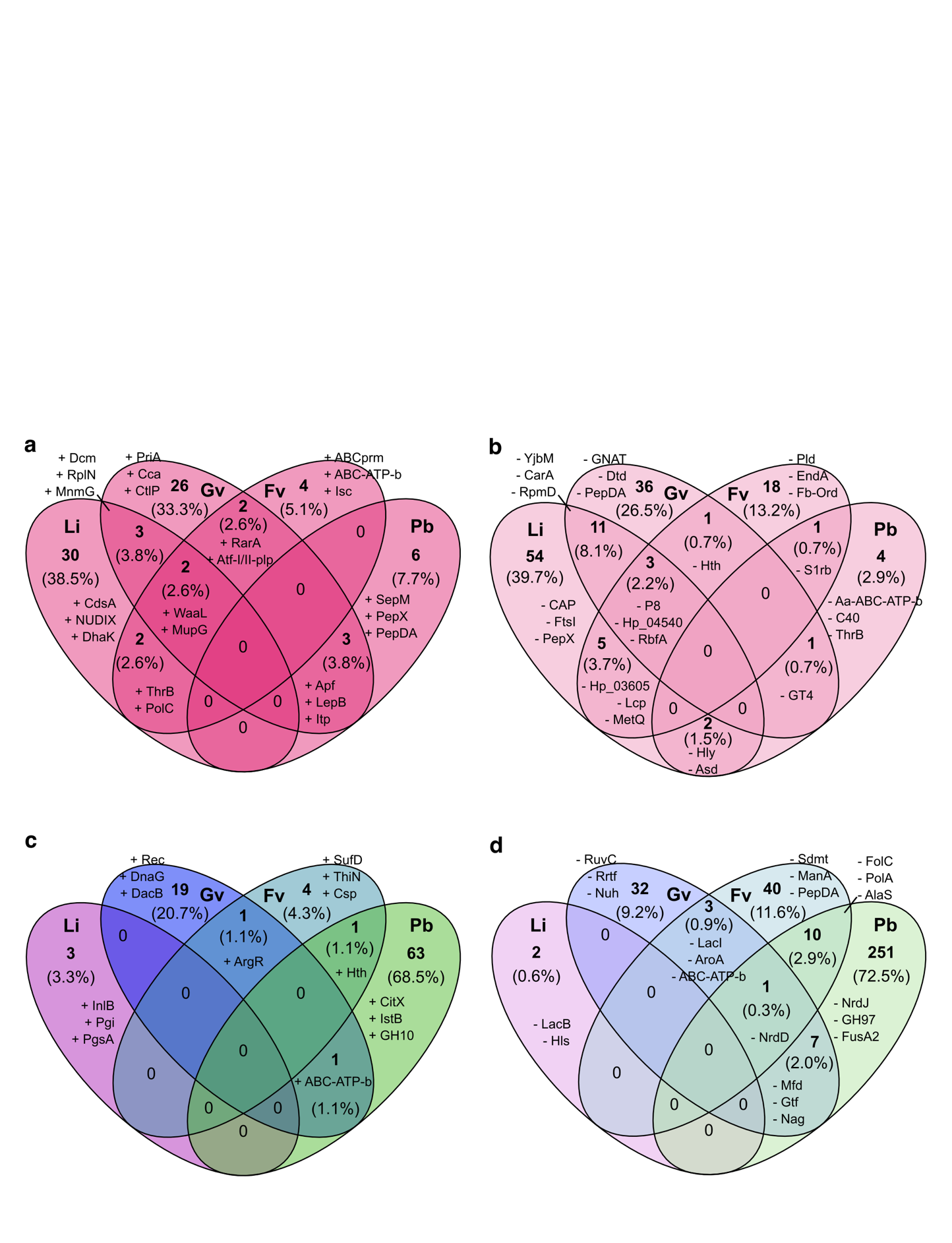

**Supplementary figure 3.** **Venn diagrams depicting unique and shared differentially expressed proteins across vaginal bacterial species following *Lc* co-culture*,* including responses in *Lc*.** a) Venn diagram showing number and relative percentage of upregulated proteins in *Lc* when coculturing with *Li, Gv, Fv* and *Pb*. b) Venn diagram showing number and relative percentage of downregulated proteins in *Lc* when coculturing with *Li, Gv, Fv* and *Pb*. c) Venn diagram showing number and relative percentage of upregulated proteins in *Gv, Fv* and *Pb* in co-cultures with *Lc*. d) Venn diagram showing number and relative percentage of downregulated proteins in *Li, Gv, Fv* and *Pb* in co-cultures with *Lc*. Abbreviations and colours: *Lc* = *Lactobacillus crispatus* (pink), *Li* = *Lactobacillus iners*, *Gv* = *Gardnerella vaginalis,* *Fv* = *Fannyhessea vaginae* and *Pb* = *Prevotella bivia*. Dark colours = upregulation, light colours = downregulation. Full protein names for abbreviations used in the figure are provided in Supplementary Table 4.

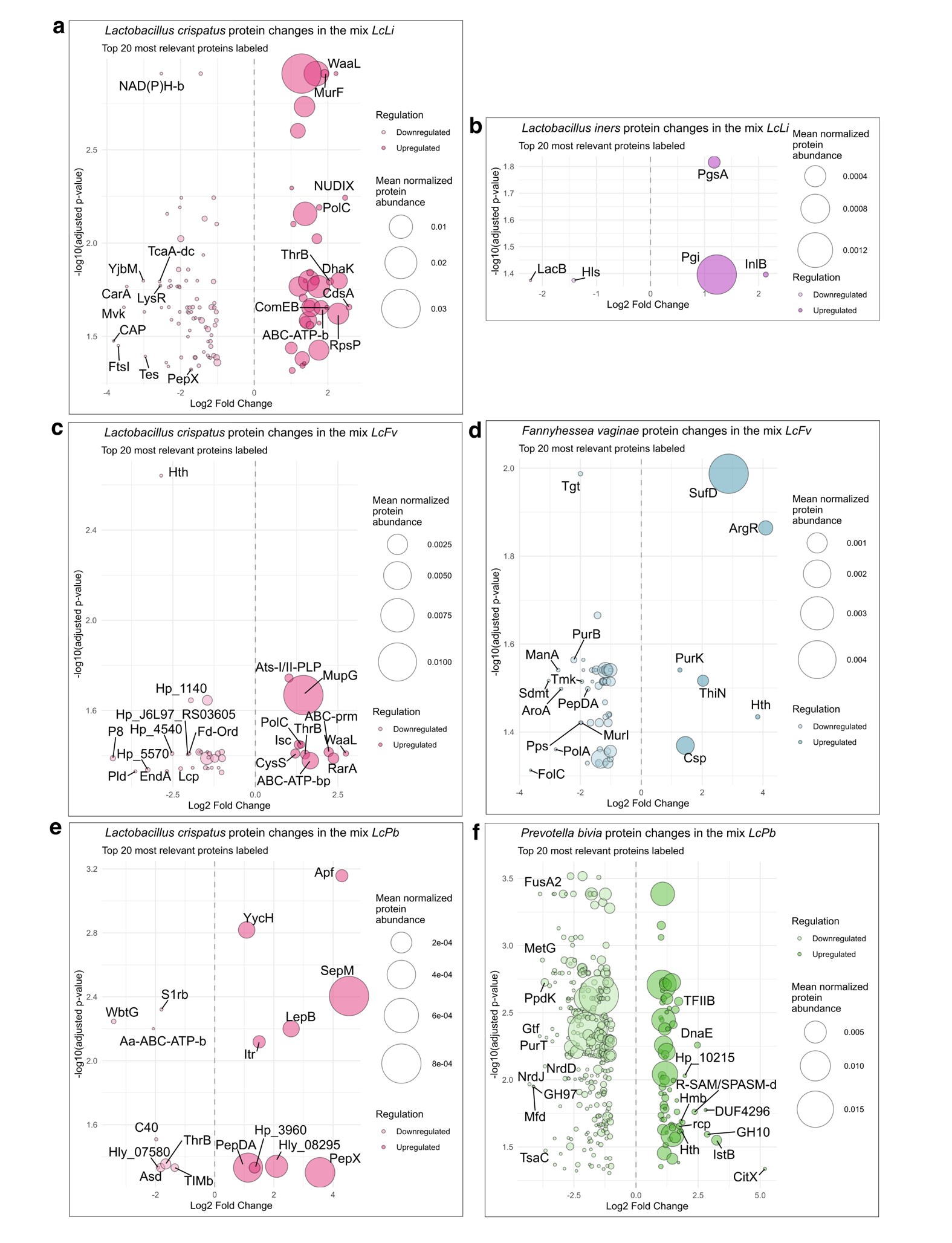

**Supplementary figure 4.** **Bubble plots of differentially expressed proteins across vaginal bacterial species following *Lc* co-culture*,* including responses in *Lc*.** a) Bubble plot showing *Lactobacillus crispatus* protein changes in the mix *LcLi*. Downregulated proteins are shown on the left side in light colour while upregulated proteins appear on the right side, Circle size indicate the mean normalized protein abundance. The top 20 proteins are labelled, prioritizing the magnitude of fold-change (|FC| > 1) over adjusted p-value. b) Bubble plot showing *Lactobacillus iners* protein changes in the mix *LcLi*. c) Bubble plot showing *Lactobacillus crispatus* protein changes in the mix *LcFv*. d) Bubble plot showing *Fannyhessea vaginae* protein changes in the mix *LcFv*. e) Bubble plot showing *Lactobacillus crispatus* protein changes in the mix *LcPb*. f) Bubble plot showing *Prevotella bivia* protein changes in the mix *LcPb*. Abbreviations and colours: *Lc* = *Lactobacillus crispatus* (pink), *Li* = *Lactobacillus iners* (burgundy), *Fv* = *Fannyhessea vaginae* (turquoise) and *Pb* = *Prevotella bivia* (green). Dark colours = upregulation, light colours = downregulation. Full protein names for abbreviations used in the figure are provided in Supplementary Table 4.

**Species-specific proteomic responses of *Lactobacillus iners* (*Li)* to other vaginal bacteria**

*Li* upregulated fructose metabolism and cell wall regulation (PfkB, ~2.92; YycH, ~2.04) when co-cultured with *Gv, Fv*, or *Pb*. In addition, increased post-transcriptional regulation (CvfB, ~1.34) with *Gv* and *Fv*, protein translation (RplV, ~8.80) with *Fv* and *Pb*, and stress protection and response to fluctuating conditions (YitT, ~1.85; TypA, ~1.86) with *Gv* and *Pb* were observed in *Li* after co-culture. Species-specific upregulated responses included mucin-binding proteins with *Lc* (InlB, 2.13), methionine transport with *Gv* (M-ABC-ATP-b, 2.61), ATP independent energy-efficient transport with *Fv* (MFS, 2.36), and protein translation with *Pb* (RbfA, 6.40) (Supplementary Fig. 5a, 6a, 6c, 6b and Figure 3g). These findings indicate that *Li* primarily enhances membrane transportation and stress response as a defense mechanism against other species.

Common downregulation in *Li* included protein translation and oxidative stress response with *Gv, Fv* and *Pb* (RplQ, ~-4.41; NfOrs, ~-3.64), quality control of nucleotide pool with *Gv* and *Fv* (X-/dl-TP, ~-3.80), membrane protein and translation quality control with *Fv* and *Pb* (HtpX, ~-3.79; TrmD, ~-2.13), and prophage transition control with *Gv* and *Pb* (Ant, ~-2.62). Partner-specific downregulations in *Li* were: lactose and galactose processing (LacB, -2.22) with *Lc*, protein translation (RplL, -4.73) with *Gv*, response to stimulus (Shk, -2.53) with *Fv*, and cell membrane assembly (AcpP, ~-4.02) with *Pb* (Supplementary Fig. 5b, 6a, 6c, 6b and Figure 3g). Thus, diverse biomolecules quality control and response to stimulus were the major downregulated pathways in *Li* when it encountered other BV-associated species.

Co-culture with *Li* induced a shared upregulation of the response to environmental fluctuations (TypA, ~2.61) in *Lc* and *Fv*, protein translation in *Lc* and *Pb* (RpsJ, ~1.64). In addition, unique protein responses after interaction with *Li* induced cell membrane maintenance in *Lc* (CdsA, 2.57), adaptive immunity in *Gv* (Cas6e, 2.45), cell wall membrane synthesis in *Fv* (IspE, 4.33), and nutrient acquisition and structural integrity in *Pb* (ExbD/TolR, 1.88) (Supplementary Fig. 5c, 6b, 6d, 3a, and Figure 3g). Overall, *Li* promoted shared stress and translation responses, while inducing species-specific adaptations in membrane/cell wall maintenance and nutrient acquisition.

Common downregulated proteins included entities involved in maintaining cellular structural integrity in *Lc* and *Gv* (FtsI, ~-2.41), ATP dependent transport in *Gv* and *Fv* (ABC-ATP-b, ~-2.22), protein translation in *Fv* and *Pb* (MetG, ~-4.69), anaerobic/microaerophilic energy balance maintenance in *Lc* and *Fv* (AdhE, ~-1.43), essential amino acid production and cell wall integrity in *Lc* and *Pb* (Asd, ~-1.52), and protein translation in *Gv* and *Pb* (RplL, ~-4.28). Species-specific vulnerabilities included: *Lc* downregulated cell wall maintenance (CAP, -3.82), *Gv* repressed vitamin salvage and metabolic competition (TenA, -5.48), *Fv* exhibited a collapse of cell wall homeostasis and amino acid and protein synthesis (MltG, -6.74; AsnA, -5.55, respectively), and *Pb* suppressed redox homeostasis (BtuE, -5.12) (Supplementary Fig. 5d, 6b, 6d, 3a, and Figure 3g). These protein-expression changes indicate that *Li* exposure disrupted cell homeostasis and downregulated protein translation across multiple vaginal bacteria.

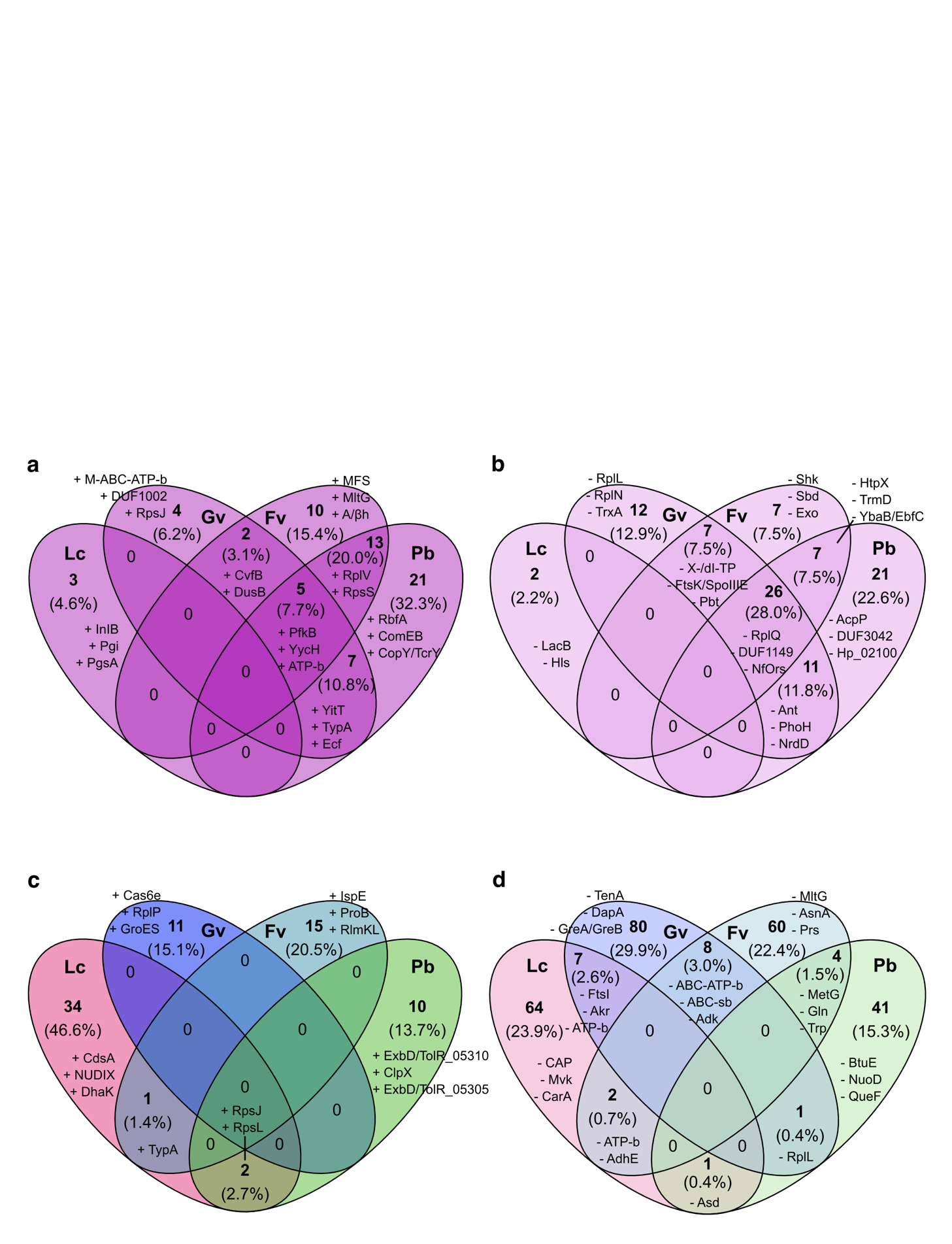

**Supplementary figure 5.** **Venn diagrams depicting unique and shared differentially expressed proteins across vaginal bacterial species following *Li* co-culture*,* including responses in *Li*.** a) Venn diagram showing number and relative percentage of upregulated proteins in *Li* when co-culturing with *Lc*, *Gv, Fv* and *Pb*. b) Venn diagram showing number and relative percentage of downregulated proteins in *Li* when coculturing with *Lc*, *Gv, Fv* and *Pb*. c) Venn diagram showing number and relative percentage of upregulated proteins in *Lc, Gv, Fv* and *Pb* in co-cultures with *Li*. d) Venn diagram showing number and relative percentage of downregulated proteins in *Lc, Gv, Fv* and *Pb* in co-cultures with *Li*. Abbreviations and colours: *Lc* = *Lactobacillus crispatus* (pink), Li = *Lactobacillus iners*, *Gv* = *Gardnerella vaginalis, Fv* = *Fannyhessea vaginae* and *Pb* = *Prevotella bivia*. Dark colours = upregulation, light colours = downregulation. Full protein names for abbreviations used in the figure are provided in Supplementary Table 4

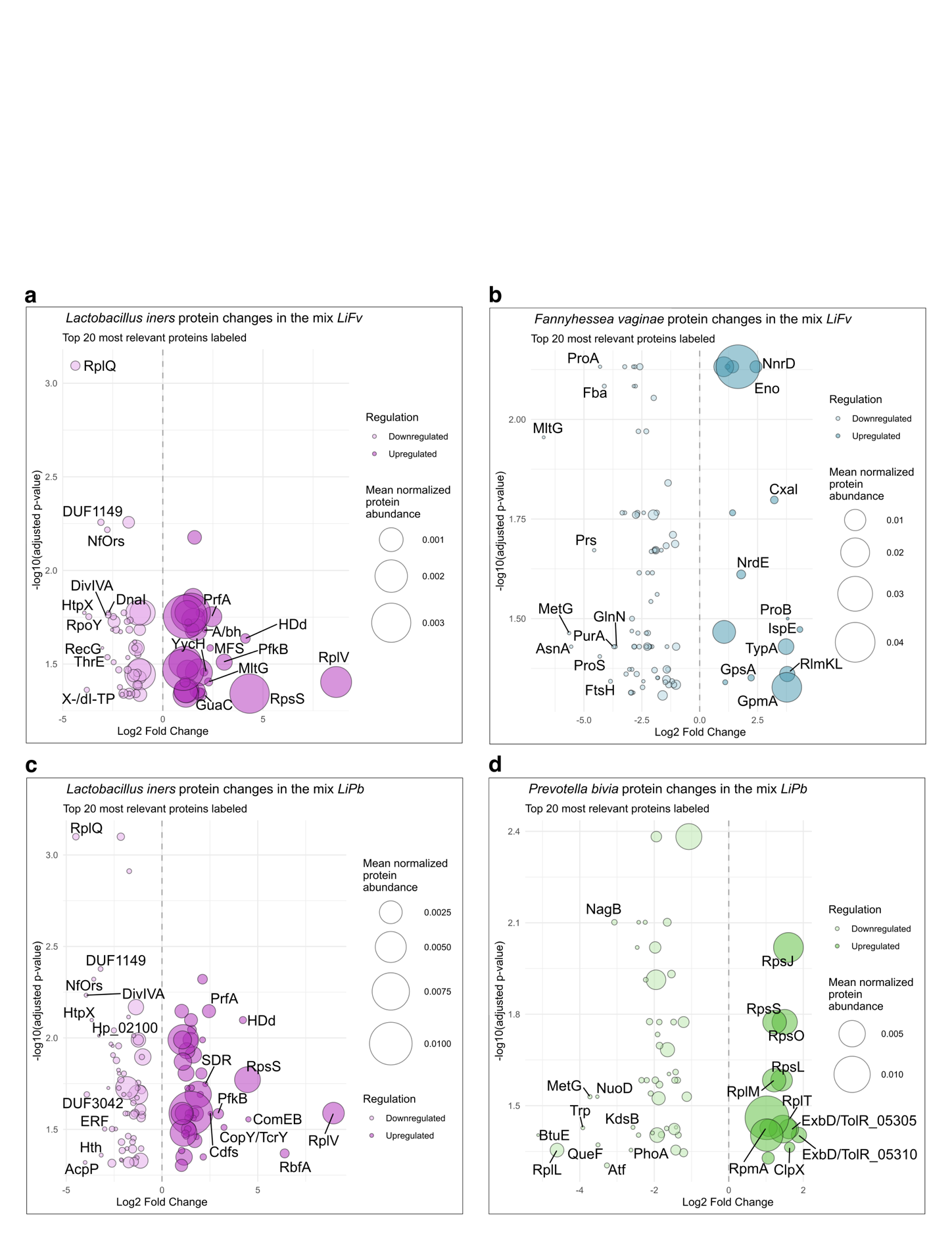

**Supplementary figure 6.** **Bubble plots of differentially expressed proteins across vaginal bacterial species following *Li* co-culture*,* including responses in *Li*.** a) Bubble plot showing *Li* protein changes in the mix *LiFv*. Downregulated proteins are shown on the left side in light colour while upregulated proteins appear on the right side, Circle size indicate the mean normalized protein abundance. The top 20 most relevant proteins are labelled, prioritizing the magnitude of fold-change (|FC| > 1) over adjusted *p-value*. b) Bubble plot showing *Fv* protein changes in the mix *LiFv*. c) Bubble plot showing *Li* protein changes in the mix *LiPb*. d) Bubble plot showing *Pb* protein changes in the mix *LiPb*. Full protein names for abbreviations used in the figure are provided in Supplementary Table 4.
